# Reanalysis of in vivo drug synergy validation study rules out synergy in most cases

**DOI:** 10.1101/2020.11.13.378430

**Authors:** Olaf van Tellingen, Renee X. de Menezes

**Author notes:** Corresponding author: Olaf van Tellingen; Division of Pharmacology, room H3.014; Plesmanlaan 121, 1066 CX, Amsterdam, The Netherlands.

## Abstract

In a recent paper in Nature Communications, Narayan et al ^1^ described the development of an *in silico* method to predict synergy of drug combinations, validated *in vivo* using 5 different models for 5 different drug pairs. The published analysis of the *in vivo* experimental data used the Chou and Talalay Combination Index, which ignores important aspects of the study design, including, but not limited to the variability between individual mice and the longitudinal nature of the data. Furthermore, the original publication reported the Combination Index as static numbers without properly accounting for experimental variability. When 95% confidence intervals are recalculated using bootstrapping methods, 4 out of 5 drug pairs do not show a statistically significant synergistic effect. Three models (originally figures 5A, B and E) are subject to severe inaccuracies in the data handling and reporting. In one model, (originally figure 5D), the data better supports an additive effect between drugs rather than synergy. As the Chou and Talalay Combination Index is poorly suited to this data, we applied mixed-effects models fitted to the longitudinal tumor growth data as an alternative means of estimating synergy. The results support the findings from the bootstrapping analysis, namely that the majority of the drug pairs do not achieve a synergistic effect. We conclude that the *in vivo* validation of the *in silico* method to predict synergy of drug combinations as proposed by Narayan et al has a negative outcome when analyzed with appropriate statistical methods.

## Introduction

Demonstrating the impact of a given drug using *in vivo* models is challenging because effect sizes are often small^2^. Because the intrinsic variability in the outgrowth of tumors *in vivo* is relatively high, achieving adequate statistical power requires sufficiently sized groups. As the complexity of the experiment increases, so too does the minimum adequate group size. A statistically sound distinction between additive and synergistic drug effects requires even larger datasets, as do comparisons between multiple combinations of drugs.

The theoretical foundation of estimating drug interaction *in vitro* by the methods as described by Chou and Talalay is well established ^3^. The basis of this method is the Combination Index (CI), where a CI<1 implicates synergy, C=1 additivity and CI>1 antagonism. Importantly, the calculation of the CI requires testing of each individual drug and their combination in a fixed ratio at multiple dose levels to create dose effect curves and assess IC_50_ values ^4^. Chou has also applied this methodology to demonstrate synergy *in vivo* using 65 mice divided over 13 groups of 5 mice ^5^. In this setup, each drug alone and its combination were tested at 4 different dose levels together with one untreated control group. Notably, 5 mice per group sufficed as the variability within groups was modest. Narayan et al ^1^, however, have used a shortcut by testing single agents and the combination at just 1 dose level, disobeying the requirement to create dose-effect curves.. Moreover, in some cases, the selected dose level for a single drug treatment was at the extreme end of the response curve. For example, in one model, docetaxel alone already resulted in 98.4% tumor volume reduction. Adding osimertinib and AZD2014 have limited impact relative to the base reduction in tumor volume achieved by docetaxel: *i.e*. an additional 0.9%, resulting in a final reduction of 99.3%. Although this marginal increase resulted in a calculated CI of 0.55, indicating synergy, the relevance of this number is questionable. Osimertinib and AZD2014 each alone reduced tumor growth by 71% and 88%, respectively, further casting aspersions on the synergistic potential of these drugs as a combination therapy.

Based on this methodology, the authors claim that the *in silico* predicted synergy was confirmed in 5 models, thereby validating the usefulness of their *in silico* strategy. Considering the aforementioned criticisms of the study power and statistical methodology for demonstrating synergy, we chose to conduct an independent analysis of the *in vivo* data. First, we recalculated the 95% confidence intervals of the CI using a bootstrapping approach, which, unlike the original publication, takes into account the intrinsic variability of these *in vivo* experiments. Although calculating confidence intervals via bootstrapping represents an improvement, it is important to recognize that Chou and Talalay’s Combination Index does not adequately capture longitudinal developments. In the original paper, the response at each tumor measurement event are handled as discrete data points, independent of previous and subsequent measurements. We have built a mixed-effects model that takes into account the course of the tumor growth per mouse, thus respecting the longitudinal nature of these data.

## Methods

Upon request, we received the raw data files of the last senior author (Bart Westerman) consisting of an excel file containing the bioluminescence imaging (BLI) data and of 5 Graphpad Prism files containing the information on survival. Later during this investigation, we received an updated version of the excel file. All files are available as supplementary data.

### Confidence intervals for the Combination Index

Chou and Talalay’s CI was originally proposed for use with *in vitro*-produced data for drug sensitivity at various doses for the drug combinations in fixed ratios. With a similar experimental design using multiple dose levels and sufficiently sized groups, it can also be applied to *in vivo* data ^3^. Narayan et al ^1^ used relatively small sample sizes, without multiple dosages for the drug and/or its combinations. This means that, in this case, the CI cannot adequately model drug interaction, as the physicochemical massaction law is not fully measured. The CI as calculated in Narayan et al. can only be applied simplistically, and will be highly sensitive to variation between individuals. Interindividual variation can be considerable due to the nature of the animal models used.

To take these issues into account, we recalculate the CI per time point as in Narayan et al. using stratified bootstrapping to better reflect experimental variability ^6^. The stratification is used per treatment (control, drug 1, drug 2 or both drugs), and ensures that resampling considers the group structure. For details, see the supplementary file “recalculations_withBootstrap_MixedEffModels.pdf”

### Models to estimate drug synergy from longitudinal data

The drug combination studies performed by Narayan et al ^1^ collected data for various time points, for the same individual mice. Analyzing data per time point ignores this aspect, and typically has less power to find effects than if data for all available data points were used. Indeed, the natural correlation between measurements for the same individual across time points yields more accurate and robust estimates, by virtue of correcting for random variation observed in individual data points.

Here, we use a mixed-effects model ^7^ approach to analyze the data, per drug pair. This allowed us to model a random effect for each individual mouse. For each drug pair, we fitted two models: one with only the main effect for each drug, which here we will refer to as the *additive* model, and one including an interaction between the two drugs, which we will refer to as the *interaction* model. We first check that the additive model yields an acceptable fit of the data. Subsequently, we test if the interaction effect is statistically different from 0. If the drugs have no interaction effects, the data may be well described by the additive model and the interaction parameter estimate from the interaction model will not be significantly different from 0. If, however, the interaction model represents a significant improvement compared to the additive one, this means that the combined effect of these drugs cannot be explained by additive effects only, so drug interaction must be occurring. For details about the models fitted see the suppl. file “recalculations_withBootstrap_MixedEffModels.pdf”.

Mixed-effects models are often used to study treatment effect on tumor growth rates ^2,8^, and can be fitted with widely available statistical software.

### Overall survival

For completeness, we also studied treatment effect on overall survival. Specifically, we compared a model with only additive effects to one which included treatment interaction, using Cox-proportional hazards models. Model fits made use of robust variance estimates ^9^.

### Software

For all analyses, we used R v 3.6.3 and packages boot v 1.3-25 (Angelo Canty and Brian Ripley; 2020; boot: Bootstrap R (S-Plus) Functions). R package version 1.3-25), lme4 v 1.1-23 ^10^ and survival 3.2-7 (Therneau T; 2020; A Package for Survival Analysis in R. R package version 3.2-7, URL: https://CRAN.R-project.org/package=survival).

## Results

Narayan et al ^1^ claim the successful *in vivo* validation of synergy in 5 separate experiments, which they present in Figure 5. Experiments 5A and B concern a U87 glioblastoma model, 5C a triple negative breast cancer model, 5D a melanoma model and 5E a chronic myeloid leukemia model. Growth kinetics of the tumors was assessed by weekly BLI measurements, which were used to calculate CI values at each measurement event. Next to that, recorded overall survival times were reported as Kaplan Meier curves.

### Experiments 5A, 5B and 5E

We first conducted a gross examination of the data. Apart from methodological concerns, we discovered a number of issues in data handling and reporting that are not in line with good scientific practice. A detailed report of our findings is available in the supplementary information (Detailed Analysis.doc). To summarize the most striking examples:

- Results from two animal experiments (5A and 5B) were pooled, although these experiments had a different study design and were conducted at different institutions. Moreover, this was not disclosed in the methods.
- The survival curve of experiment 5A was not based on overall survival, but on a highly arbitrary date of progression assessment. This was also not disclosed in the methods.
- The p-values associated with survival analyses were artificially inflated via the use of a one-sided t-test, rather than a log-rank test, as is the norm in the field. The actual calculations contained many flaws, such as comparing mean versus median values, and the use of censored animals to increase the sample size and improve p values.
- Depicted p values for survival involved the comparison of the combination versus the control group only, neglecting the comparison between the co-administered drugs and single drugs. This is not clearly stated in the methods section, and is does not capture synergy in any way.

Aside from these issues, there are additional concerns regarding the experimental protocol and mistakes that affected the results. Concerning experiment 5A:

- There were many faulty injections of luciferin, in some cases also affecting measurements at day 0, which is the reference for all subsequent BLI measurements.
- Instead of using the ratio of the BLI data of each individual mouse relative to its own BLI taken at day 0, the tumor growth was calculated by taking the ratio of the group mean BLI values of each day relative to the group mean BLI value on day 0.
- The last measured BLI data of animals that died during the course of this study were used as input values for the remaining days (carry last value forward). Effectively, this means that low BLI values of animals that died in the triple combination group due to toxicity affected (reduced) the group mean values on subsequent days, whereas it is likely that these tumors would have progressed at some point in time, had the animal survived the treatment.
- All calculations were performed on linear data, whereas a log-transformation of BLI data should have been applied.

After addressing these issues (see supplementary file Detailed Analysis.doc for details), the drug combination in experiment 5A was no longer synergistic (Fig. 1). In fact, docetaxel was already highly efficacious when given alone; adding the other drugs did not result in further improvement.

**Figure 1.**
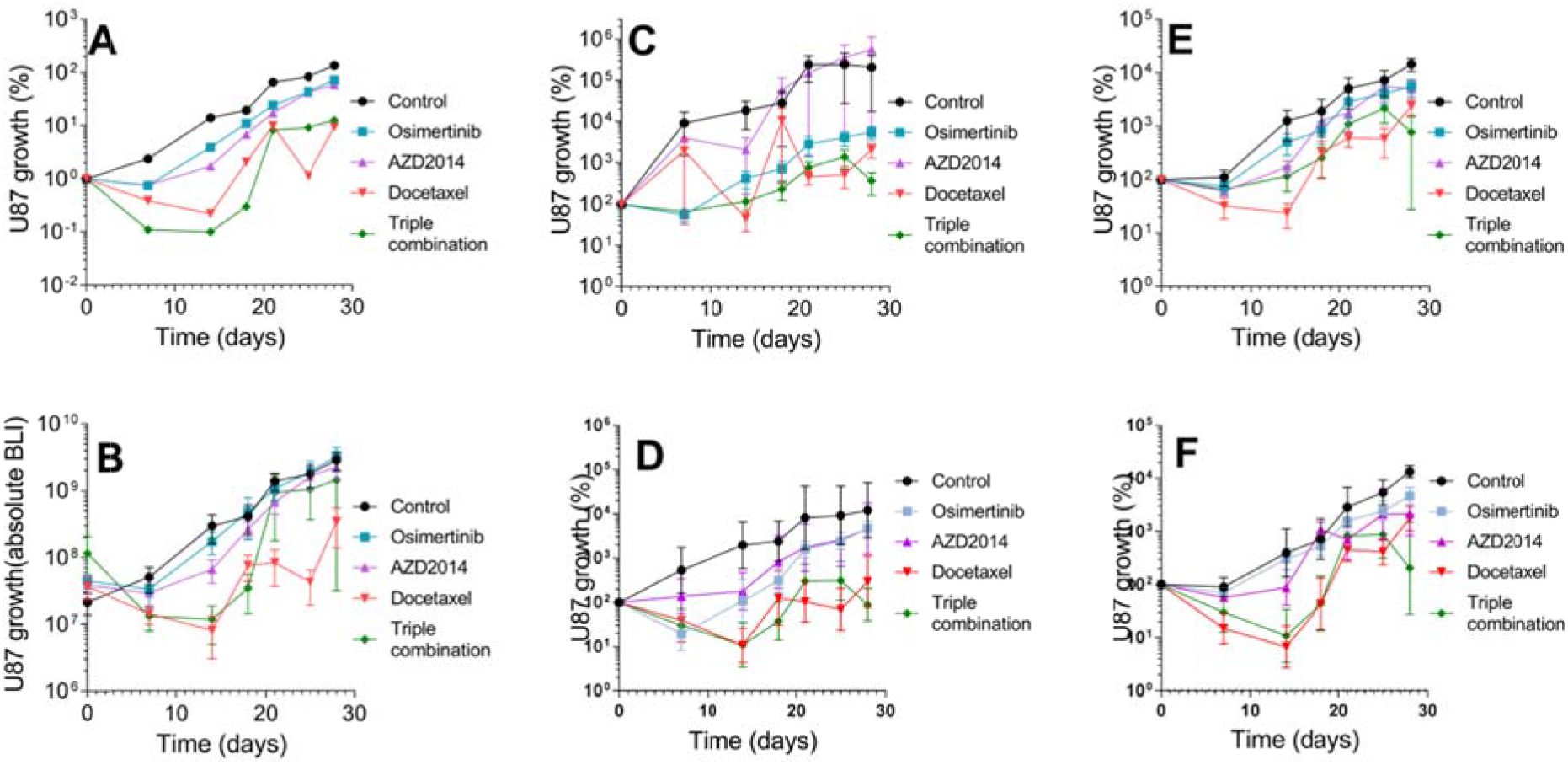
Re-Analyzed data of Experiment 5A. **A)** Data as used in the paper. Growth(%) was calculated from group means BLI per day normalized to the group means at day 0 (Start of treatment). **B)** Same original data depicted as mean absolute BLI values (± SE) without normalization. **C)** Tumor growth(%) calculated as BLI values of each animal relative to its own BLI value at start day. **E)**: The same data after rejection of incorrect injections and carry last value forward data points. Panels **D** and **F** are similar to **C** and **E**, respectively, but after log conversion of the BLI data. In panels B-E, standard errors were computed each time point separately ignoring other time points.

Similarly, the drug combination in experiment 5B was not synergistic. The experiment was underpowered with only 4 animals per group, and the actual CI determination involved even fewer animals due to early dropouts.

In the case of experiment 5E, the original synergistic outcome was due to an error (reference to wrong cells) in the excel file. After correction, the CI values are close to 1; therefore, additive drug effects cannot be ruled out. The survival plots support the conclusion that the effect is additive. Both drugs are about equally active, increasing the median survival from 35.5 (control) to 58 (imatinib; +23 days) and 60.5 (dasatinib + 25 days), and to 75 days (+40 days) with the combination.

For these reasons, we did not analyze the data for 5A, 5B and 5E further, considering only the remaining 2 experiments in the remainder of this document.

### Experiments 5C and 5D

When looking at the tumor growth curves and survival plots, experiment 5C most clearly stands out as a possible example of synergy (Fig. 2a). The response on AZ628 alone has an almost negligible effect, but when given with gemcitabine it considerably augments the efficacy relative to gemcitabine alone. Of course, this is the result of the comparison at one dose level only, but the CI values are very low (<0.1), indicating a large effect size. Indeed, experimental variation is relatively low and 95% bootstrap confidence intervals for the CIs comfortably reject the hypothesis of no synergy (Fig. 2b). In contrast, the outcome is much less clear for experiment 5D (Fig. 2a). Both CGP-082996 and gemcitabine alone reduce tumor growth to 57% and 32% at day 28 relative to untreated controls, respectively. In contrast, the combination reduces growth to 16%, resulting in a CI of 0.61 for that day. However, due to large experimental variation, the confidence intervals for the CI include values close to 1 (Fig. 2b), suggesting that the reduction in tumor growth is not sufficient to exclude purely additive effects.

**Figure 2.**
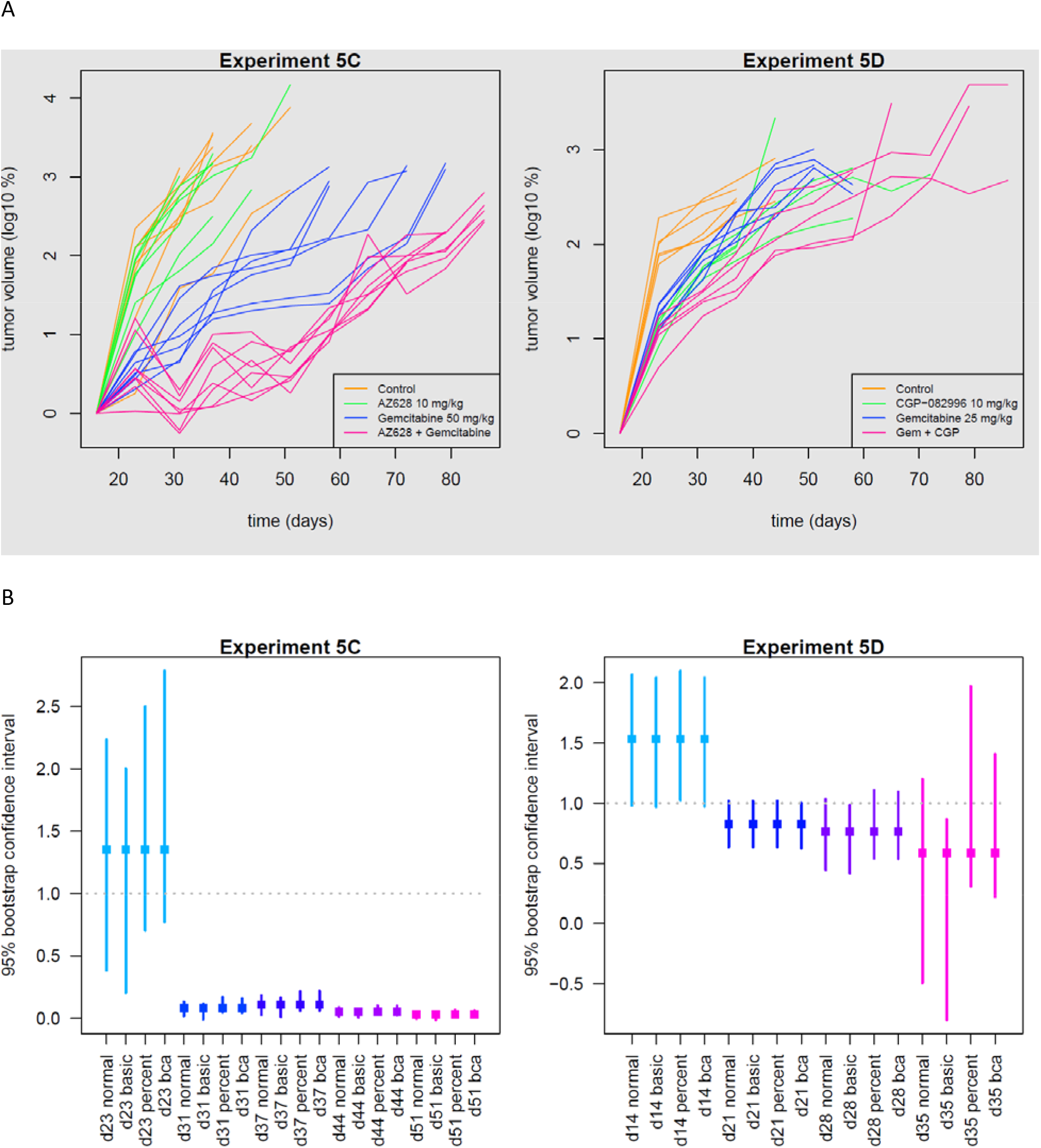
A. Growth curves of individual mice, depicted as the ratio of BLI values relative to BLI on the start day of treatment for experiment 5C (left) and 5D (right). B. Bootstrap analyses to determine the 95% confidence interval of the calculated combination indices.

As an alternative to the Chou and Talalay methodology, a mixed-effects model was fitted to the data for all time points. Synergy was determined by comparing two models: one fitted with only additive effects of the two drugs, and one that also included an interaction effect between both drugs. These comparisons confirmed what was suggested by the combination index’s confidence intervals: a synergistic effect for figure 5C, and an additive effect for figure 5D. These conclusions were in agreement with survival analysis of the overall survival.

In summary, 4 of the 5 *in vivo* validations of synergy are negative when proper statistical tests are employed.

Details about the analyses and all results can be found in the supplementary file “recalculations_withBootstrap_MixedEffModels.pdf”

## Discussion

In the work of Narayan et al ^1^, Chou and Talalay’s Combination Index was used to evaluate combined drug effects in a suboptimal way that led to incorrect conclusions. This was partly due to inadequate experimental design, which meant that the Combination Index failed to fully quantify dose-effect relations. In addition, large experimental variation and small sample sizes meant that extreme values had an unduly large impact on estimates. We illustrated this by computing confidence intervals for individual Combination Indices. We here propose using a mixed-effects model using the course of tumor growth per mouse to compensate for individual variation and to better capture longitudinal effects.

A fresh look at the experimental data of Figure 5 of Narayan et al. determines that synergistic findings for experiments 5A, 5B and 5E were due to insufficient sample size, problems with the experimental methodology, and errors in calculations. Reanalysis of data for 5C and 5D which takes all time points into account confirmed synergy for 5C only. This aligns with conclusions drawn from confidence intervals for individual CI values, as well as from overall survival analysis.

It is a well-known fact that, when studying treatment effects from repeated tumor measurements, models which consider individual time points sequentially yield less power than those that consider all time points collectively ^8^. It is therefore to be expected that the same holds true when testing for treatment interaction effects. Our results suggest this to be the case.

We conclude that the claims of synergy as predicted by the *in silico* strategy presented by Narayan et al ^1^ cannot be confirmed in 4 out of 5 validation experiments. This was partly due to severe flaws in data handling and partly due to the use of statistical methodology inappropriate for the experimental design. Alternative methods presented here lead to robust conclusions across multiple approaches.

## Supporting information

Detailed findings of manuscript review

Document with calculations in R

Graphpad Prism file Figure 1 reanalysis

Datafolder containing original data files Bart Westerman

## Notes

**Conflict of Interest statements**: OvT: None RM was recently recruited as group leader of the Biostatistics Centre of the Netherlands Cancer Institute. Before this she was a biostatistician at the VUmc, which is the institute of Narayan and colleagues. RM is a coauthor of the paper under discussion, but she was not involved in the analyses of the *in vivo* data.

https://www.nature.com/articles/s41467-020-16735-2

### Competing Interest Statement

OvT: None
RM was recently recruited as group leader of the Biostatistics Centre of the Netherlands Cancer Institute. Before this she was a biostatistician at the VUmc, which is the institute of Narayan and colleagues. RM is a coauthor of the paper under discussion, but she was not involved in the analyses of the in vivo data.

