## Supplementary material for "Reanalysis of in vivo drug synergy validation study rules out synergy in most cases": Detailed findings of manuscript review

A detailed description of the issues in data handling and reporting found in the paper of Narayan et al.

Bioluminescence imaging (BLI) measurements were used as a surrogate for the amount of tumor in the body of the animal. We first received the BLI data of the study presented in figure 5A in the paper. This study was conducted at the Amsterdam UMC, whereas the other studies (depicted in 5B–E) were conducted at Harvard Medical School.

The following issues in the analysis of Fig 5A were discovered:

1. The growth was presented by taking the ratio of the group mean BLI values of each day relative to the group mean BLI value on day 0. This is unusual as BLI measurement provides longitudinal data per mouse and it is common to use the ratio of BLI data of each individual mouse relative to its own BLI taken at day 0.

2. The BLI data of animals that died in the course of the study were used as values in the remaining days (carry last value forward). Effectively, this means that low BLI values of animals that died in the triple combination group due to toxicity affect (reduce) the mean value on subsequent days, whereas it is likely that these tumors would have progressed at some point in time, when the animal would have survived the treatment

3. There are many incorrect injections of luciferin. This can be recognized by the fact that there is a sharp decrease in BLI in between two adjacent measurements. Unfortunately, this has also affected some measurements at day 0, which is the reference point for assessing the changes in tumor mass.

4. The data are presented in Fig 5A on a log-scale, which is correct since tumor progression is a logarithmic process, but all calculations are done with linearized data. However, BLI data can have a wide range, especially in this case with the many incorrect luciferin injections. For 5A the range on start day of treatment varies from 1.33x10^5^ to 6.61x10^8^, thus more than three orders of magnitude. When using linearized data, the highest values in a set weighs much more heavily in the calculation of the group mean than lower values. For example: the mean value of the observations 1, 3, 5, 7, 9, 1000 is about 170. By using log-conversion, this issue is largely repaired (the mean of the above mentioned observations will be 9.9). Due to the fact that linear data instead of log-converted data were used, the synergy as calculated in this series is completely dependent on one high value BLI (animal 7R) on the day of randomization in the triple combi group (Excel file OvT1: field I33: 6.61x10^8^). This value is by far the highest of all animals at the day of randomization. If this value is changed to 6.61E7, the calculated values for the combination index will become (see below):

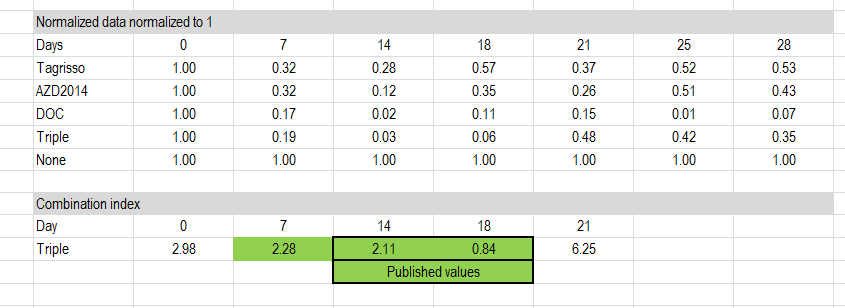

Source: Excel file Copy of Data_request+OvT2_Synergy_complete_with raw IVIS data_after_correspondance2 v7/7/2020

The fact that changing one data point can have such a dramatic impact, demonstrates the very limited robustness of determining synergy using linear data.

To visualize the results, the data of 5A was analyzed in several ways.

Panel A shows the data as used in the paper, group mean BLI per day normalized to the group mean at start day.

Panel D shows the same original data as mean BLI but now without normalization to the start day.

Panel B shows tumor growth as BLI data of each animal relative to BLI at start day.

Panel C: shows the same data after deletion of incorrect injections and last value forward data points.

Panels E and F are similar to B and C, respectively, using log conversion of the BLI data.

Error bars indicate SE.

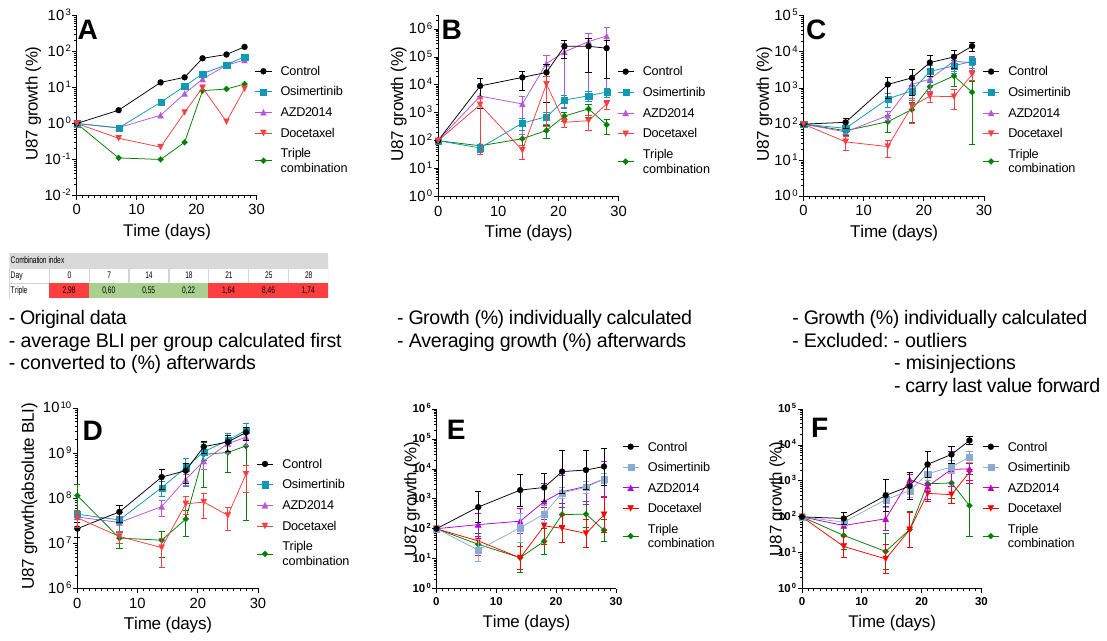
(source: Graphpad prism file: Narayan et al NC reanalysis Fig 5a_OvT; embedded. Please double click to open, works only when Graphpad Prism is installed on your computer)

Conclusion: It is obvious that in all cases, except panel A, the triple combination is not better than docetaxel alone.

In case of Fig 5B. It appears that this analysis is based on a very small set of animals.

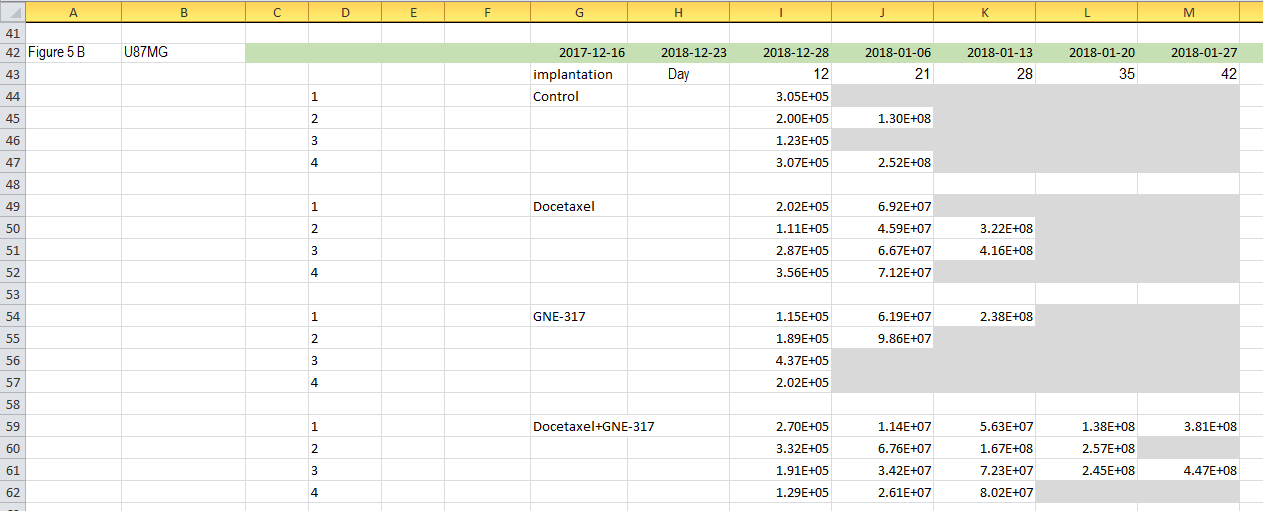

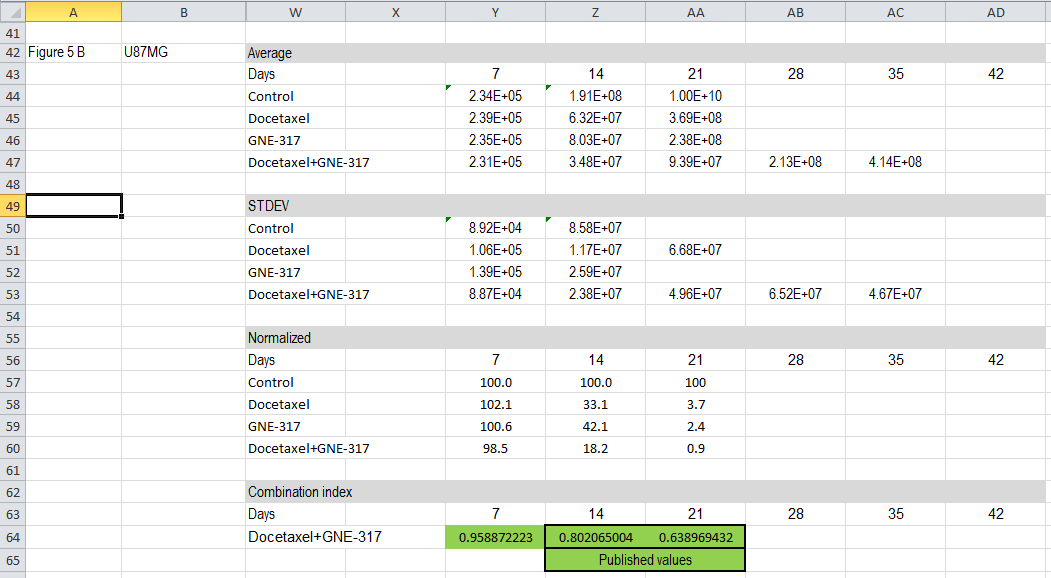

Source: Excel file Copy of Data_request+OvT2_Synergy_complete_with raw IVIS data_after_correspondance2 v7/7/2020

Each cohort starts with 4 animals, but already at day 21 (column J in the top excel sheet) several animals have died. At day 28 all control animals are lost, effectively making it impossible to calculate the combination index. In order to repair this issue a value of 1x10^10^ is has been typed in as surrogate (AA41 in bottom excel sheet). Other things that stand out are:

There are no BLI values taken at the day of randomization (start of treatment).

The days as presented (7, 14, 21; column Y, Z, AA) are not in line with the actual day (columns: I, J, K, L). If treatment was started at day 7 (as described in the methods) then day 12 (2018-12-28) is actually day 5 (not 7; column Y). On the other hand, it may also be that day 12 is actually the day of randomization, since the mean BLI values of the 4 cohorts are similar, whereas in the previous experiment (5A), treatment with docetaxel resulted in a clear response at day 7. In conclusion, this experiment is under-powered and contains too few observations to justify any claims of synergy.

Next, we go to the analysis of the results in Fig 5E.

In an earlier version of the excel file that I obtained from the corresponding author, I noted a mistake in the excel spreadsheet (reference to wrong cells). Instead of using the Normalized data for entry into the formula for the combination index, the Average BLI data was used.

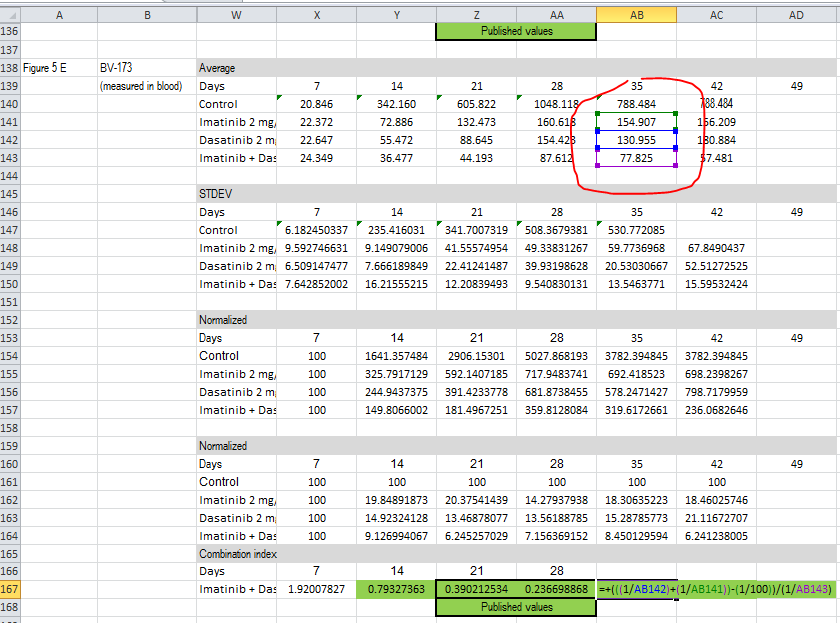

The corresponding author acknowledged that this is a mistake and has sent a corrected version.

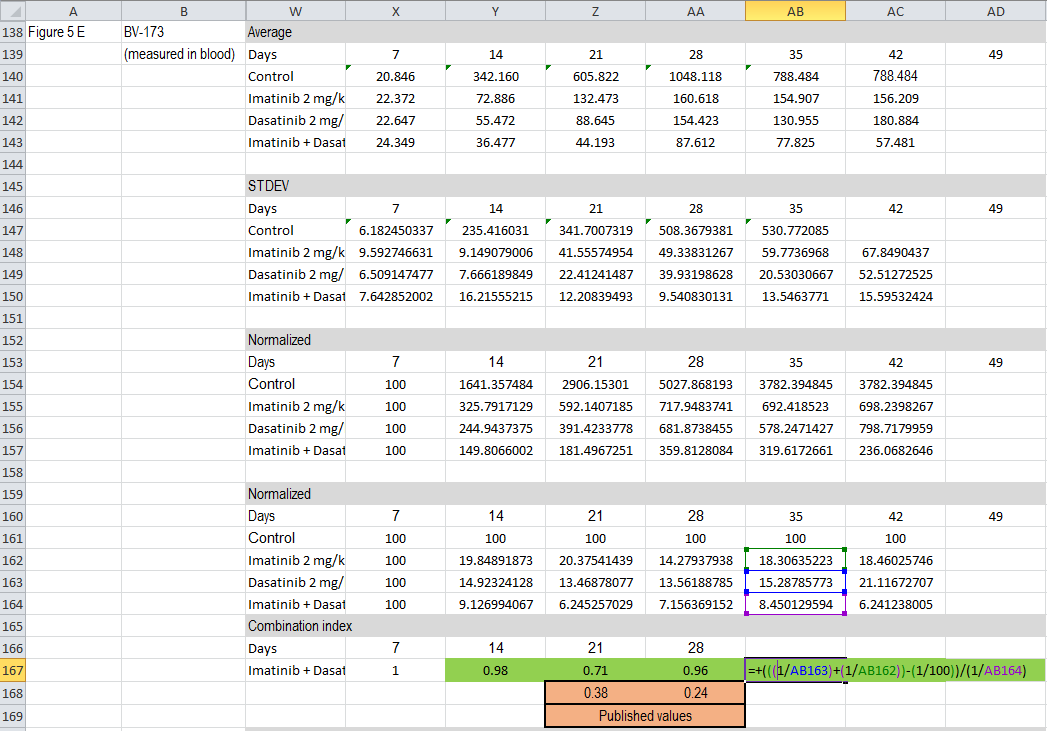

Source: Excel file Copy of Data_request+OvT2_Synergy_complete_with raw IVIS data_after_correspondance2 v7/7/2020

Only at day 21, the combination index is below 0.80, whereas it is around 1 at all other days.

The significance of this value of 0.71 on one day in the series will be addressed further in the analysis, but on all other days the combination index is very close to 1, indicating absence of synergy.

The excel data files for experiments 5D and 5E appear to be correct. Notably, however, Figure 5D contains an error. The growth percentages of CGP-082996 and the combination at day 28 are 32% and 16%, respectively, instead of 3% and 0.3%.

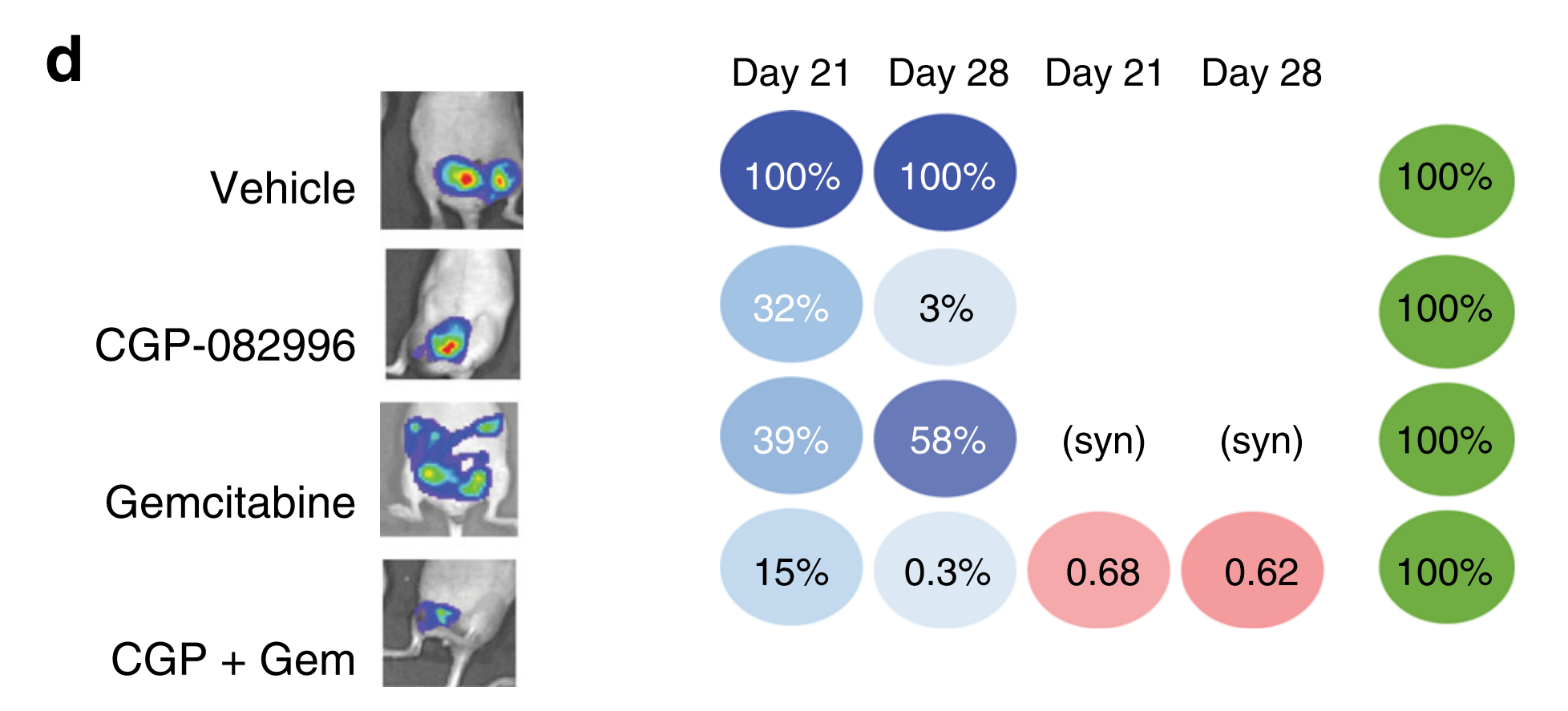

Source: Figure 5 of the manuscript.

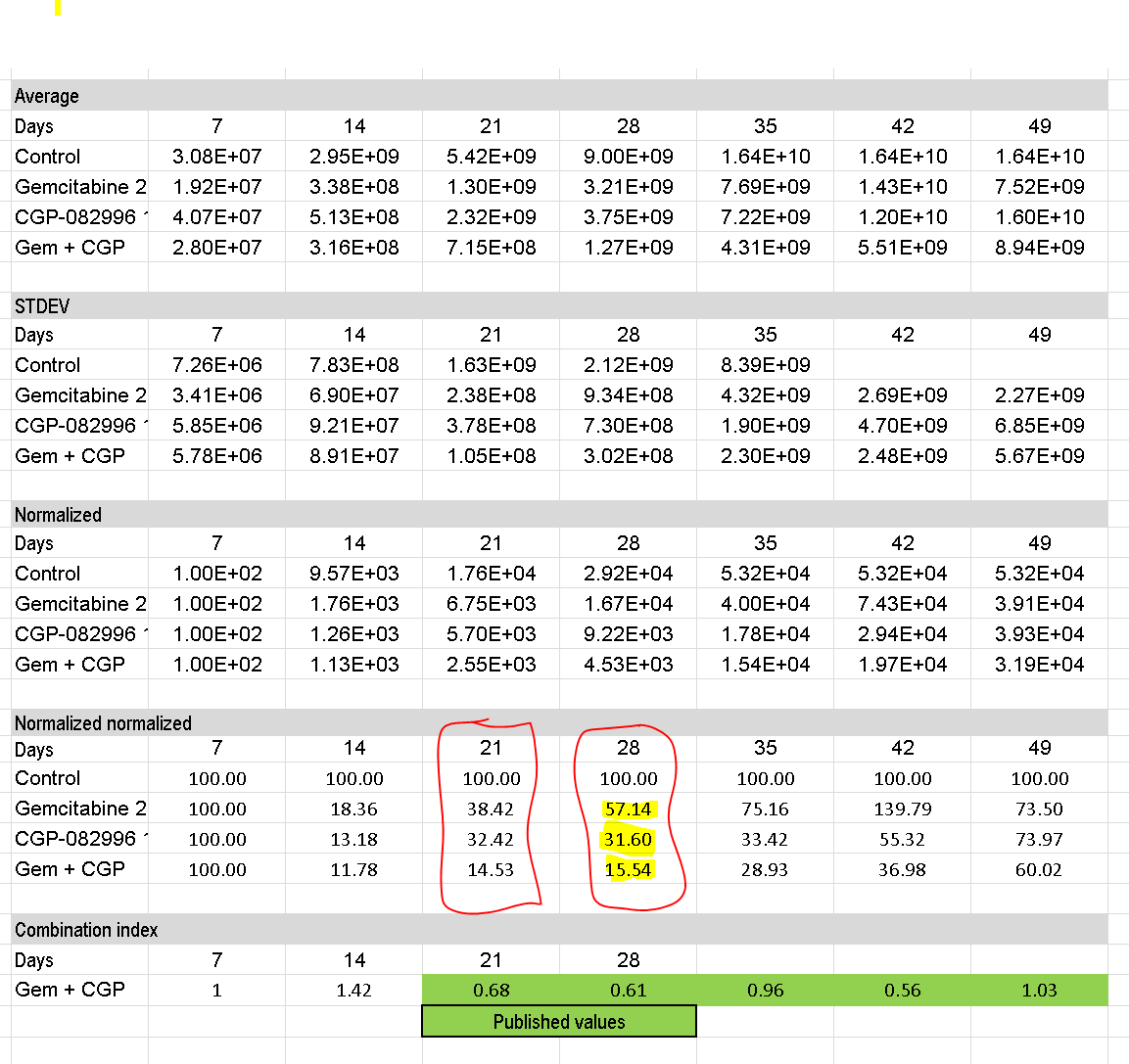

Source: Excel file Copy of Data_request+OvT2_Synergy_complete_with raw IVIS data_after_correspondance2 v7/7/2020

So far for now for the results of the BLI data measurements. Besides the calculation of tumor growth by BLI the paper also provided data on the survival. Below is what is written in the legend of Fig. 5 (The red lines refer to survival results):

***f Kaplan–Meier curves showing a better survival of mice treated with the combination of drugs (see Supplementary Data 5). For all experiments, luciferase levels were normalized to levels of one week after injection. Toxicity monitoring consisted of assessment of body weight, hematopoietic-, liver-, and brain toxicity. P-value: t test (one-sided) of the median survival. The number of mice per group are shown in the figures.***

There are several issues:

Without access to the raw data files it is unclear that the p values that are given in the survival plots actually comprise the comparison between the combination groups *versus* the control group. So, actually the curves do not show a better survival of mice treated with the combination of drugs. They merely show survival of the combination versus the control groups. This is not clear from the description in the legend and there is no justification why this has been done. It does not seem very logical to confirm synergy by only testing the control group versus the combination group. It is also not reasonable to expect that the readers will actually realize that this has been done.

The calculation of survival was not done by a standard methodology, such as Log-rank test, but was done by a t-test comparing the day of death of the control group animal versus the combination group. For undefined reasons, sometimes the median value was used, sometimes the mean. Notably, the t-test was only one-sided and there was no correction for multiple comparisons. Again, this information is not easily accessible to the reader.

When checking the analysis of Experiment 5A, a discrepancy between the number of animals used for plotting the survival curves and the animals in the study (see GrapPad file Figure 5A Survival) was noted. There were 7 animals in the osimertinib (=Tagrisso), AZD2014 and Triple combi, but 10 in the control group and 11 in the docetaxel. The number of animals in Experiment 5A were actually 7 for all treatment groups and 6 for the control group. The corresponding author explained that the extra animals in the control and docetaxel cohorts were ‘borrowed’ from experiment 5B. Quoted from the senior author: “In the field, it is sometimes applied to combine experiments with mice to increase power, which was done here. Experiment 5A and B have been combined to improve power”. However, it is obvious that mixing the results from two experiments, conducted at different moments and in different labs (Amsterdam and Boston) is incorrect and unjustified. Another point was that there were BLI values recorded from animals on days after the recorded day of death in the survival analysis. The corresponding author explained that the survival analysis of experiment 5A was not done on the day of death of the animals but on the day of progressive disease, since the endpoint could not be determined because many mice got sick from toxicity. This was not clear from the methods. In the methods there is a statement that: “Progressive disease is defined as the last time point before disease progression (i.e., weight loss)”. Weight data, however, were not provided and it is not clarified how body weight loss due to toxicity was discerned from weight loss due to tumor progression. Based on the BLI data in the spreadsheet, a graph of the BLI data was prepared in order to visualize how time to progression relates to the course of the BLI signal in three of the five cohorts (Control, docetaxel and triple combination groups).

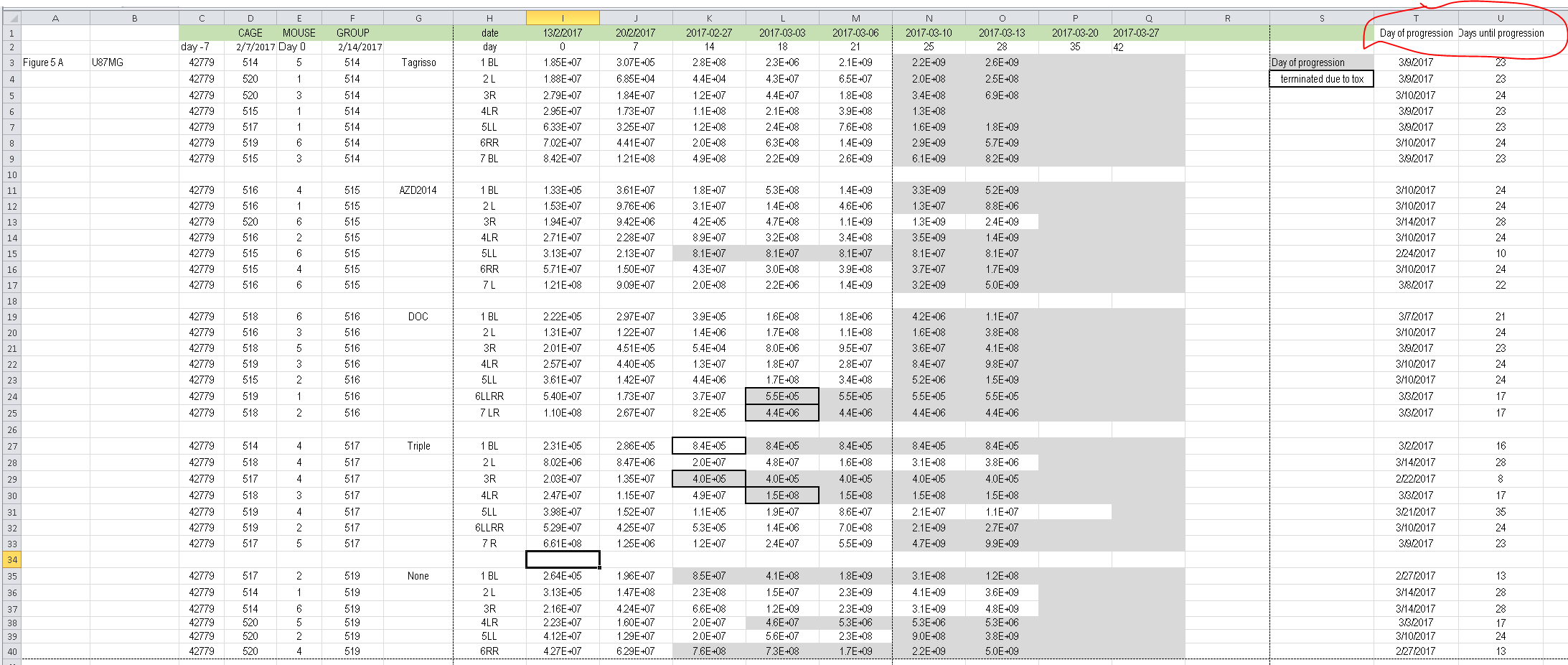
Excel file showing tumor growth (BLI-data) and presumed date of progression in columns T and U.

Source: Excel file Copy of Data_request+OvT2_Synergy_complete_with raw IVIS data_after_correspondance2 v7/7/2020

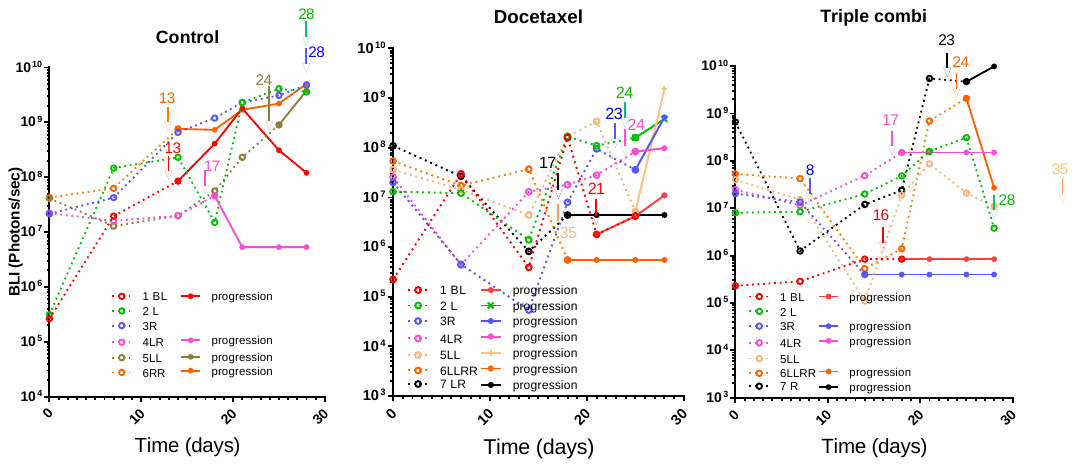
Graphical presentation of the tumor growth curves of individual mice of Control, docetaxel and triple combi group. The time of tumor progression for each animals is indicated by the arrow. From that point onwards the dotted line becomes continuous and the symbols closed. (Source: Graphpad Prism file: embedded, please double click to open, works only when Graphpad is installed on your computer)

It is clear from this graphical presentation that there is no clear justification for the chosen day for “time to progression”. Some animal experiencing weight loss still survived for (several) weeks, while others still had very small tumors when progressing.

Although it seems clear that the “survival” data of experiment 5A are unfit to be used for a survival analysis, the data was used to reproduce the survival analysis using the methodology in the paper (one-sided t-test).

The survival analysis that was actually presented in the paper with this yielded a p = 0.0078 (Control vs Triple combi group). Several ‘mistakes’ had to be made in order to reproduce to this number.

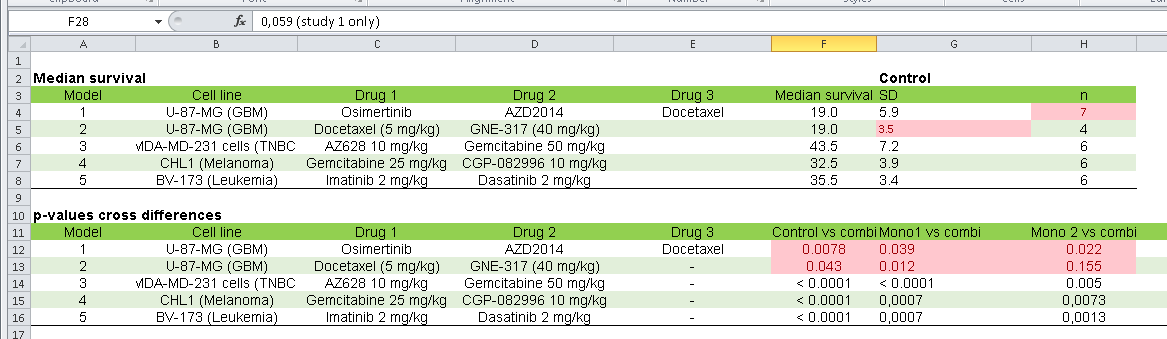
Source: excel file: Supplementary Data 5_annotated_OvT.xls (This file was taken from the Nature Comm. Website: Suppl info)

Below is an excel sheet containing the data of this experiment.
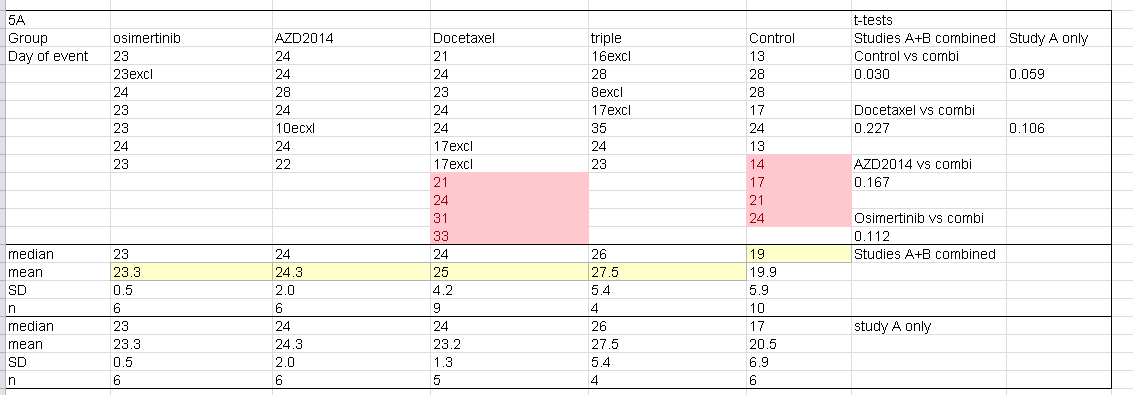
Source: excel file: Supplementary Data 5_annotated_OvT.xls

The following numbers had to be used in Graphpad (see below) to reproduce the data.

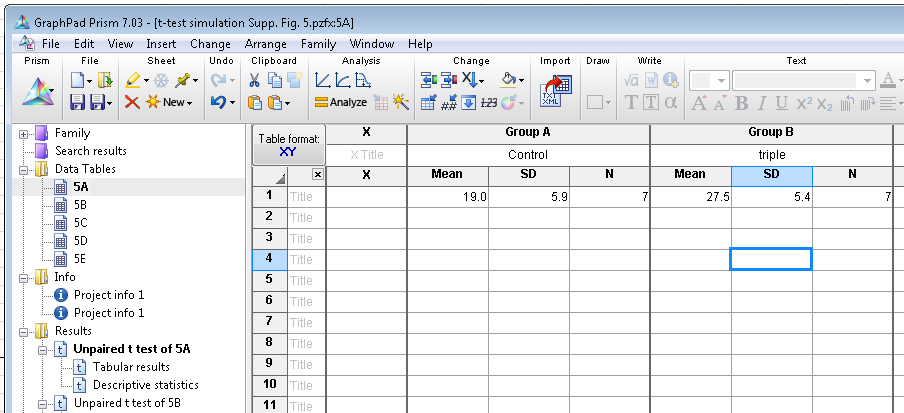

Output:

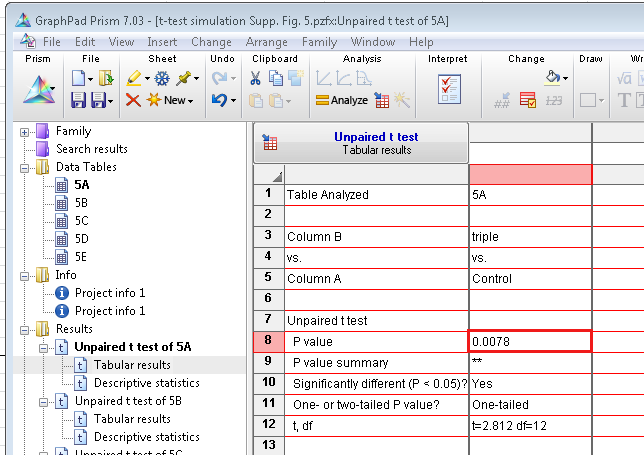

What was not correct is:

-The mean of the triple is compared to the median of the control. However, the median cannot be used in the (parametric) t-test. Note that the mean was 19.9 and the median 19.0. Thus using the median, makes the difference bigger in this case.

-The number of observations in the triple group is actually 4 instead of 7 since 3 mice were censored. Yet, 7 is being used as input, which again improves the p-value.

-The number of observations in the Control group is 6 (or 10 with the 4 ‘borrowed’ mice). In this (wrong) analysis using 10 would have been ‘logical’, but would have revealed the use of more animals. However, 7 is also incorrect.

Although as mentioned above the “time-to-progression” data is unsuited, we conducted an survival analysis using the data of Experiment 5A only (leaving out data of experiment 5B):

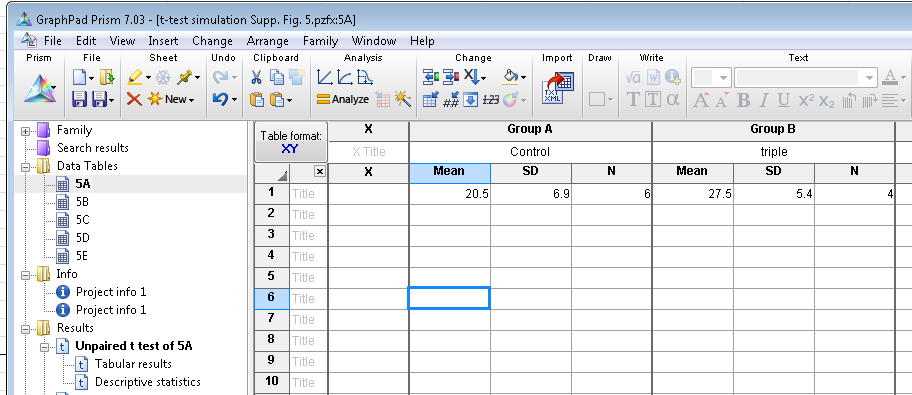

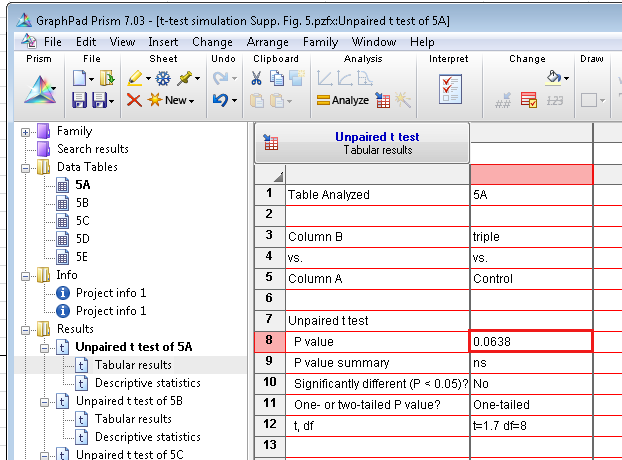

This analysis returns a p value of 0.0638, even under the most lenient conditions (one-sided, no multiple comparison correction).

Also in the other experiments, there were several inaccuracies (indicated in pink).
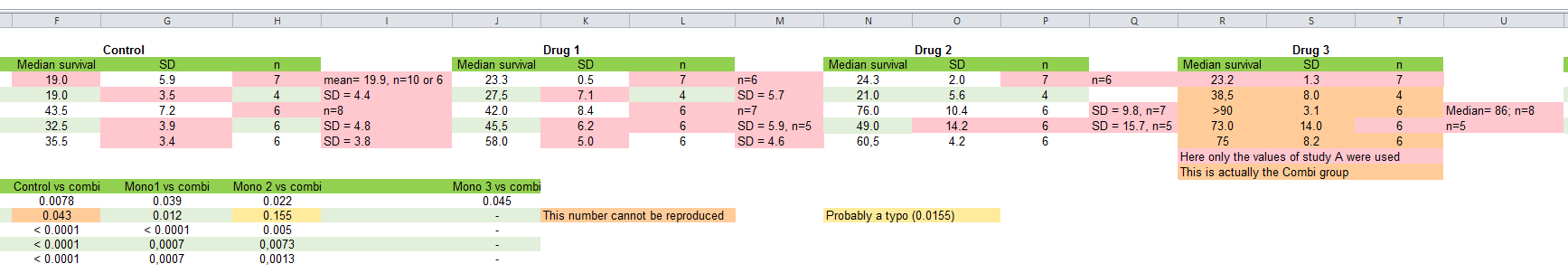
 Source: excel file: Supplementary Data 5_annotated_OvT.xls

These, however, did not change the significance of the outcome when comparing control group versus the combination treatment. But this is caused by the lenient conditions under which the t-tests were conducted.

Re-analyses of the survival using the Log-rank (Mantel Cox) was conducted using Graphpad Prism files as provided by the senior author. Data of Figure 5E had to be corrected, since this file actually contained a copy of the data from experiment 5D.

Exp. 5B.

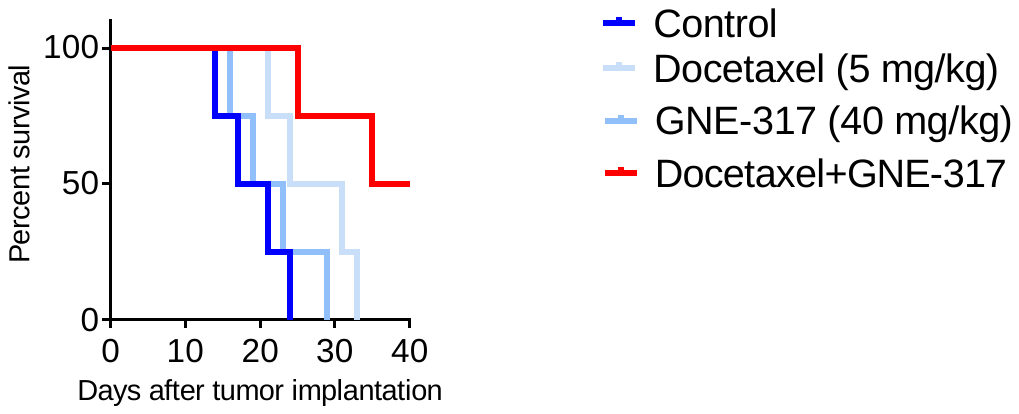

The Graphpad Prism file of survival analysis returns a p value of 0.0249 in the overall group comparison, meaning that there are significant differences between any of these curves. In a post-hoc analyses we have compared  docetaxel+GNE-317 versus  docetaxel and found a p=0.051. Note: this value needs further correction for multiple comparison (multiply by the SQRT(6)=2.45 (4 groups can be compared in 6 ways). Thus the corrected for multiple comparisons p-value = 0.125. This means that the combination is not significantly better than docetaxel alone. Notably, this is a grossly underpowered study.

Exp 5C.

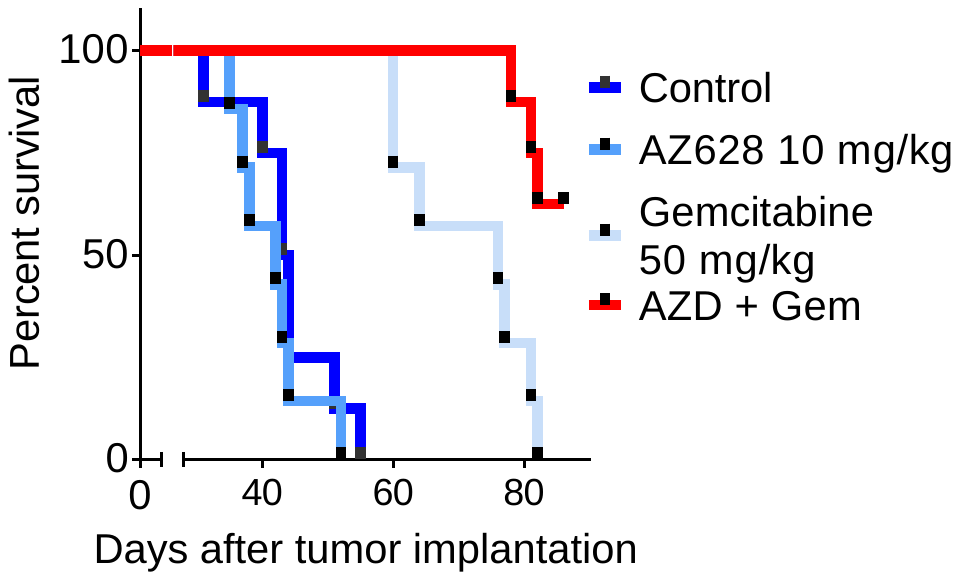

The Graphpad Prism file of survival analysis returns a p value of <0.0001 in the overall group comparison, meaning that there are significant differences between any of these curves. In a post-hoc analyses we have compared  the combi versus gemcitabine and found a p=0.0024. Following correction for multiple comparison: p=0.0058. Thus significance is reached. Since the effect of the other drug (AZD628) alone is negligible, synergy is probable.

Exp 5D.

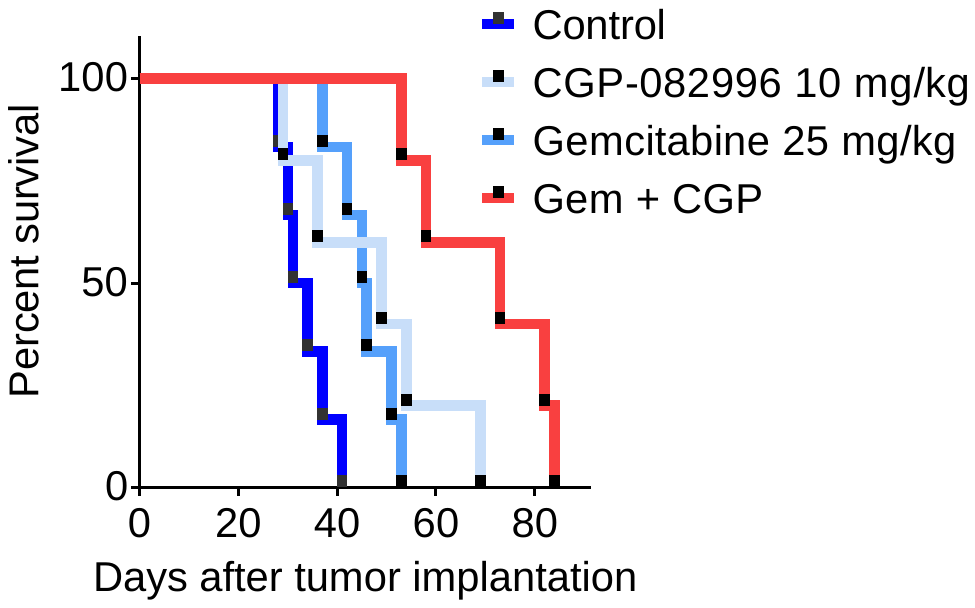

The Graphpad Prism file of survival analysis returns a p value  of 0.0009 in the overall group comparison, meaning that there are significant differences between any of these curves. Since both drugs show activity, we have compared all curves:

| Comparison |  | Multiple comparison | Significance |
| --- | --- | --- | --- |
| Gemcitabine vs CGP | 0.73 | 1.00 | NS |
| Control vs gemcitabine | 0.0022 | 0.0053 | * |
| Control vs CGP | 0.0704 | 0.17 | NS |
| Control vs combi | 0.0014 | 0.00343 | * |
| Gemcitabine vs combi | 0.003 | 0.0073 | * |
| CGP vs combi | 0.0333 | 0.081 | NS |

This means that also here significance is not reached between the combination and the single agent.

Exp 5E

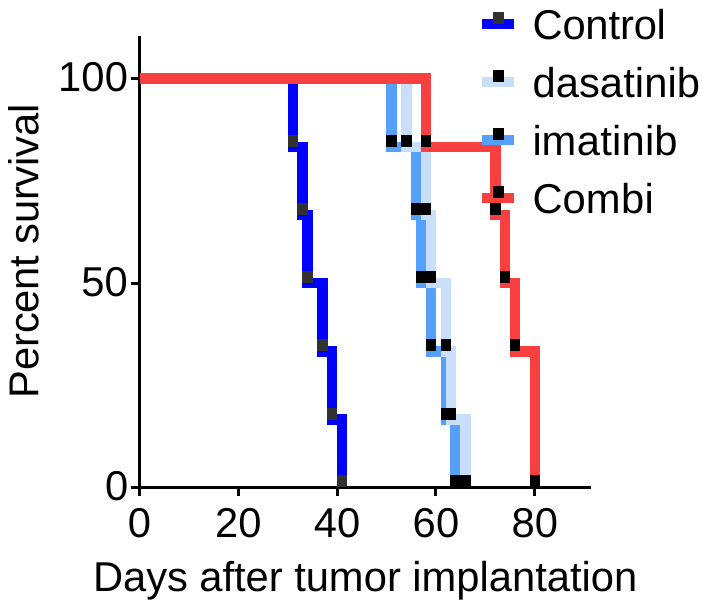

The Graphpad Prism file of survival analysis returns a p value  of 0.0009 in the overall group comparison, meaning that there are significant differences between any of these curves. Since both drugs show activity, we have compared all curves:

| Comparison |  | Multiple comparison | Significance |
| --- | --- | --- | --- |
| Imatinib vs dasatinib | 0.43 | 1.00 | NS |
| Control vs imatinib | 0.0005 | 0.0012 | * |
| Control vs dasatinib | 0.0005 | 0.0012 | * |
| Control vs combi | 0.0005 | 0.0012 | * |
| imatinib vs combi | 0.0049 | 0.012 | * |
| Dasatinib vs combi | 0.0071 | 0.017 | * |

Both drugs are about equally active, increasing the median survival from 35.5 to 58 (imatinib; +23 days) and 60.5 (dasatinib + 25 days). The combination seems additive as the survival increases to 75 days (+40 days).
