## Supplementary material for "Reanalysis of in vivo drug synergy validation study rules out synergy in most cases": Document with calculations in R

### Evaluating drug synergy

Renee Menezes

2020-10-15

### Introduction

#### General information

This document refers to recalculations of results presented in

https://www.nature.com/articles/s41467-020-16735-2

The re-analysis of data for the mouse studies in the paper will be done, considering one drug pair at a time. We refer to the drug pairs by their labels in figure 5, from `A` to `E`. This document focuses on the analysis of drug pairs `C` and `D`, but the scripts used can be easily applied to data for the other drug pairs.

For each drug pair, tumor volume is measured for individual mice at fixed time points. This yields multiple tumor volume measurements per mouse. Per drug pair, a different endpoint is used. The data involves both repeated measurements as well as time to event. Per drug pair, each mouse is assigned to a different treatment group, namely: drug 1, drug2, both drugs 1 and 2, and no treatment (control).

Tumor growth is measured on an exponential scale, so a log-transformation is used for visualization, and a logarithmic link function is used in regression models.

#### Drug synergy and the Combination Index

##### Background

The Combination Index (CI) was proposed by Chou and Talalay to quantify drug interaction. It is based on the physicochemical principles of the mass-action law, and it relies on multiple dose-response measurements available for each individual drug and for their combination, the latter in fixed ratios. The references to the method from the paper itself are:

Ref 29: Chou, T. C. & Talalay, P. A simple generalized equation for the analysis of multiple inhibitions of Michaelis-Menten kinetic systems. *J. Biol. Chem.* **252**, 6438–6442 (1977).

Ref 68: Chou, T.-C. & Talalay, P. Generalized equations for the analysis of inhibitions of Michaelis-Menten and higher-order kinetic systems with two or more mutually exclusive and nonexclusive inhibitors. *Eur. J. Biochem.* **115**, 207–216 (2005).

**Combination Index computed by Narayan et al.**

In the Narayan et al. paper, the experimental design for the data used involved a single dose-response measurement per drug pair and time point. This does not contain enough information for the CI to evaluate drug interaction as originally intended, as some parameters of the underlying dose-effect curve cannot be well quantified in the formula - see

Chou, T-C (2010). Drug combination studies and their synergy quantification using the Chou-Talalay method. *Cancer Research* **70**: 440-446. DOI: 10.1158/0008-5472.

Thus, there is no guarantee that the computed CIs in this case can adequately represent the relationship between the effects of the two drugs under study.

In addition, the sample sizes available were relatively small, relative to the inter-individual variability. Furthermore, the CIs were computed using averages of the observed tumor values per group, which does not convey variability present in the data. Yet another point is that, in some of the experiments, animals died or had only limited follow-up available, but in calculations the authors took the value at the last measured time point and projected it to later time points. This again decreases data variability (as some values are repeated across time points), and it masks effects such as treatment toxicity.

For these reasons, we decided to re-compute the CIs considering only the data available per time point. To evaluate how robust conclusions are to experimental variability, we constructed confidence intervals for the computed CIs using the bootstrap.

#### Synergy in a time-course study

The study design used by Narayan et. al. yielded tumor volume measurements per time point, for multiple pre-specified times, per treatment. By computing CI's per time point, the relationship between time points was ignored, and that may have led to loss of power to detect true effects of interaction between drugs. Indeed, by considering all measured values together, more points are available and time-point specific effects are averaged out. In order to consider all time points simultaneously, we have used a mixed-effects model.

One mixed-effects model was used per drug combination, to fit tumor volume over time per mouse. The mouse effect is random within the group, and the group is fixed. This enables us to test for a group effect, and to compare the effects of different groups. So the model we used consists of:

$$g(\mu_{ijkt}) = \alpha + b_i + \gamma_j I_j + \gamma_k I_k + \eta_{jk} I_{jk} + \delta_t,$$

where $b_i$ represents the (random) individual mouse effect with $b_i \sim \mathcal{N}(0, \sigma_i^2)$, $\gamma_j, \gamma_k$ represent individual, fixed drug effects for drugs indexed by $j, k$ on tumor volume across time (with $I_j, I_k$ binary variables indicating whether mouse $i$ received the corresponding drug or not), $\eta_{jk}$ represents the additional effect of the combination of drugs that cannot be accounted for by their additive, individual effects (with $I_{jk}$ a binary variable indicating whether mouse $i$ received both drugs or not), and $\delta_t$ represents the effect for time point $t$. This model is thus able to represent additive effects of the drugs, as well as effects that cannot be accounted for by them. A mouse which received only a single drug contributes to the estimation of the corresponding drug's effect, and a mouse receiving both drugs will contribue to estimating both individual drug effects, as well as the interaction.

The objective here is to test the hypothesis that the two drugs have additive effects, against the alternative that their effects interact, in a way that cannot be explained by the additive effects of the two drugs alone.

One thing that can be done to check if the two drugs display an interaction effect is to compare the fit of the mixed-effects model above with that of the simpler model, without the interaction:

$$g(\mu_{ijkt}) = \alpha + b_i + \gamma_j I_j + \gamma_k I_k + \delta_t.$$

For this, we first check if the effects of the individual drugs can be assumed to be additive, based on the data above. Subsequently, we add the interaction effect to the model, and check if this represents an improvement of the fit, compared with the model without the interaction.

Note that the models here proposed assume that treatment effect is constant in time. This is likely to hold as a first-order approximation. We are of course aware that a model which would include an interaction between treatment and time, representing differential tumor growth between treatments, would be more realistic. However, to fit not just the interaction between treatment effect and time, as well as the interaction between treatments, would require a considerably larger sample size, which is not available here.

##### Time-to-event analysis

We also analysed the time-to-event information, and checked if treatment had an effect on it. Specifically, we will study how time-to-event is affected by the treatment variables, with the same variables as before. In formulae, we will fit the model:

$$S(t|I_j, I_k, I_{jk}) = h(t)\exp\{\gamma_j I_j + \gamma_k I_k + \eta_{jk} I_{jk}\},$$

where $h(t)$ represents the baseline hazard.

As we did with the mixed-effects model, we will test if the parameter $\eta_{jk}$ is equal to zero. This will enable us to test if the model with additive and interaction effects represents the data better than the model with only the additive effects.

The recorded follow-up data includes a variable that represents the time at which each individual mouse reached its endpoint. This time may or may not coincide with the fixed time points at which tumor volume was evaluated. Since there is a single time-to-endpoint value per mouse, the survival model above can be fitted using classic survival models with fixed effects.

#### Data analysis

```
source("var_in_colour.R") # function to generate colours for factors
library(lme4)
library(survival)
library(boot)
```

##### Figure 5D

###### Read in and explore data

```
mydata <- read.delim("fig5d.txt")
time <- c(7, 14, 21, 28, 35, 42, 49, 56, 63, 70, 77)
```

```
mydata$group <- factor(mydata$group, levels = c("Control", "CGP-082996 10 mg/kg",
                                                "Gemcitabine 25 mg/kg", "Gem + CGP"))
```

The data read in contains information on 24 animals measured by 14 variables. Note that there are 2 animals with no tumor growth, which we now leave out of analyses.

```
mydata <- mydata[ rowSums(is.na(mydata)) < 12, ]
```

Some variables have many `NA` entries:

```
colSums(is.na(mydata))
```

```
##        ID    group       d7      d14      d21      d28      d35      d42
##         0        0        0        0        0        0        5        9
##       d49      d56      d63      d70      d77 endpoint
##        12       17       18       19       20        0
```

Visualizing the data per animal, within groups:

```
gcol <- var.in.colour(mydata$group, mystart = 0.1, myend = 0.9)
par(mfrow=c(1, 4), bg = "grey90")
for(xg in 1:nlevels(mydata$group))
{
    datag <- mydata[ mydata$group == levels(mydata$group)[ xg ], ]
    dplot <- datag[, 3:(ncol(datag)-1)]
    plot(1, 1, col = "white", xlim = range(time), ylim = range(dplot, na.rm = TRUE),
         xlab = "time (days)", ylab = "tumor volume", main = levels(mydata$group)[ xg ])
    for(xr in 1:nrow(datag)) lines(time, dplot[xr, ], col = gcol[[2]][ gcol[[3]] == levels(mydata$group)
    }
```

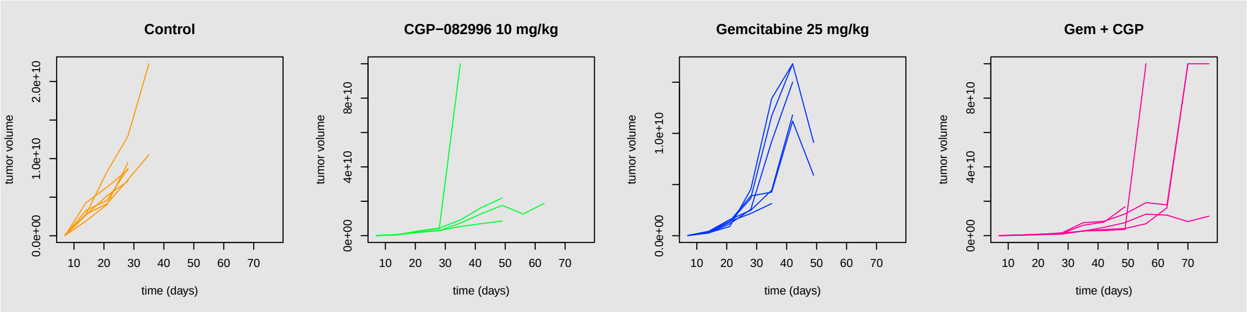

For these data, the initial value can be taken as $T = 7$. We divide the volume values by the one at $T = 7$ per mouse and log-transform the data. Then the tumor volume evolution for animals coloured by group is:

```
gcol <- var.in.colour(mydata$group, mystart = 0.1, myend = 0.9)
par(bg = "grey90")
data.all <- mydata[, 3:(ncol(mydata)-1)]
data.all <- data.all/matrix(data.all[, 1], nrow = nrow(data.all), ncol = ncol(data.all))
myylim <- range(log10(data.all), na.rm=TRUE)
plot(1, 1, col = "white", xlim = range(time), ylim = myylim,  # ylim = range(log10(dplot),na.rm = TRUE)
         xlab = "time (days)", ylab = "tumor volume (log10 %)")
for(xg in 1:nlevels(mydata$group))
{
    datag <- data.all[ mydata$group == levels(mydata$group)[ xg ], ]
    dplot <- datag
    for(xr in 1:nrow(datag)) lines(time, log10(dplot[xr, ]), col = gcol[[2]][ gcol[[3]] == levels(mydata
    }
legend("bottomright", legend = gcol[[3]], lty = "solid", col = gcol[[2]], cex = .8)
```

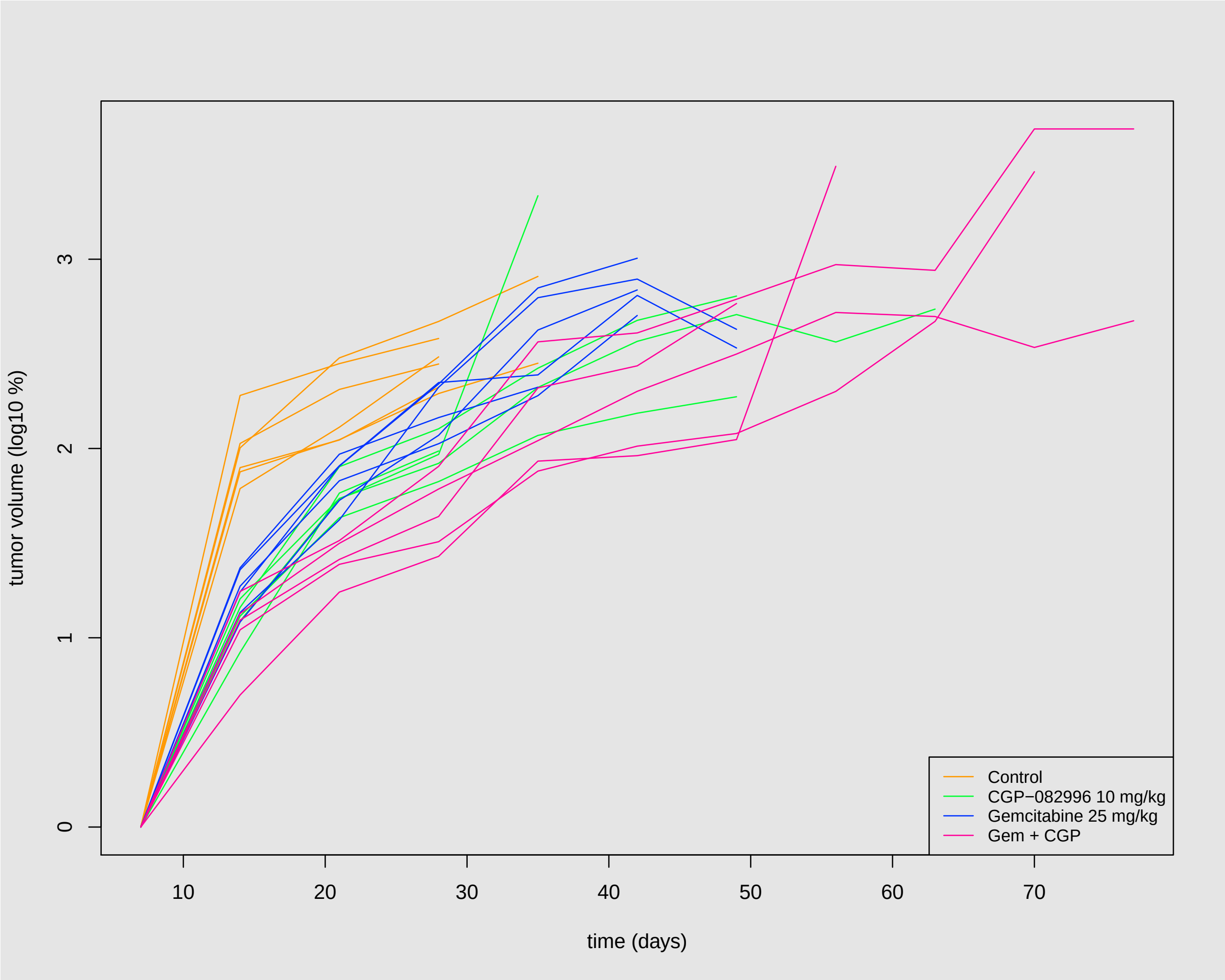

```
mydatad <- mydata
```

##### Computing the Combination Index

The combination index was computed in the original paper as follows:

- computed tumor volume average per time point and group
- divide each average by that for the initial time point
- divide each tumor volume percentage by that for the control group (group of mice which did not receive any of the drugs)

We will recompute the CIs per time point using the data for Figure 5D. Subsequently, we will construct confidence intervals for the CIs using the stratified bootstrap, which samples taking the groups into account.

```
# Function to compute combination index for all time points
# Can be used with boot - but make sure to run stratified bootstrap for multiple samples

combind <- function(mdata, xi){
  datas <- mdata[xi, 3:ncol(mdata)]
  fgr <- mdata$group[ xi ]
  datam <- apply(datas, 2, function(x) { tapply(x, INDEX = fgr, FUN = mean, na.rm = TRUE)})
```

```
  datamr <- datam/matrix(datam[, 1], nrow = nrow(datam), ncol = ncol(datam))
  datamrr <- datamr/matrix(datamr[1, ], nrow = nrow(datamr), ncol = ncol(datamr), byrow = TRUE)
  cvec <- apply(datamrr[, -1], 2, function(x) { (1/x[2] + 1/x[3] - 1/100)*x[4] } )
  cvec
}

# Define function to compute combination index per time point
# To be used with boot to run stratified bootstrap

combind.boot <- function(mdata, i){
  datas <- mdata[i, 3:ncol(mdata)]
  fgr <- mdata$group[ i ]
  # Mean per group
  datam <- apply(datas, 2, function(x) { tapply(x, INDEX = fgr, FUN = mean, na.rm = TRUE)})
  # Relative to t=7
  datamr <- datam/matrix(datam[, 1], nrow = nrow(datam), ncol = ncol(datam))
  # Relative to control
  datamrr <- datamr/matrix(datamr[1, ], nrow = nrow(datamr), ncol = ncol(datamr), byrow = TRUE)
  fcombind <- function(x) { (1/x[2] + 1/x[3] - 1/100)*x[4] }
  cvec <- apply(datamrr[, -1, drop = FALSE], 2, fcombind)
  cvec
}
```

The Combination Indices for the original data, with no data transformation, per time point, are:

```
combind.obs <- combind(mydata, xi = 1:nrow(mydata))

round(combind.obs, 3)
```

```
##      d14      d21      d28      d35      d42      d49      d56      d63
##    1.534    0.826    0.763    0.588      NaN      NaN      NaN      NaN
##      d70      d77 endpoint
##      NaN      NaN    3.175
```

Note that only the first 4 time points (discarding the initial time point, which serves as a reference) have valid results for the CIs. Indeed, for later time points all individuals in the control group were no longer available.

We now build confidence intervals for Combination Indices using the stratified bootstrap with 1000 resamples. Only CIs with valid results are used, so this is restricted to the first 4 time points. Here we use the function `boot` from the package of the same name to do this. The stratified bootstrap performs resampling within groups defined by the treatment, thereby ensuring that treatment groups are always well represented in each resample.

```
vtime <- paste0("d", time)
ntime <- 4
comb.l <- vector("list", ntime)
names(comb.l) <- vtime[2:(ntime+1)]
for(xi in 2:5)
{
  mydatat <- cbind(mydata[, 1:3],  mydata[, vtime[xi] ])
  names(mydatat)[ncol(mydatat)] <- vtime[xi]
  comb.l[[ vtime[xi] ]] <- boot(mydatat, statistic = combind.boot, R = 1000, stype = "i", strata = myda
}
```

Per time point, the empirical distribution of the indices is:

```r
for(xi in 1:4) plot(comb.l[[ xi ]]); title( paste(vtime[xi + 1]))
```

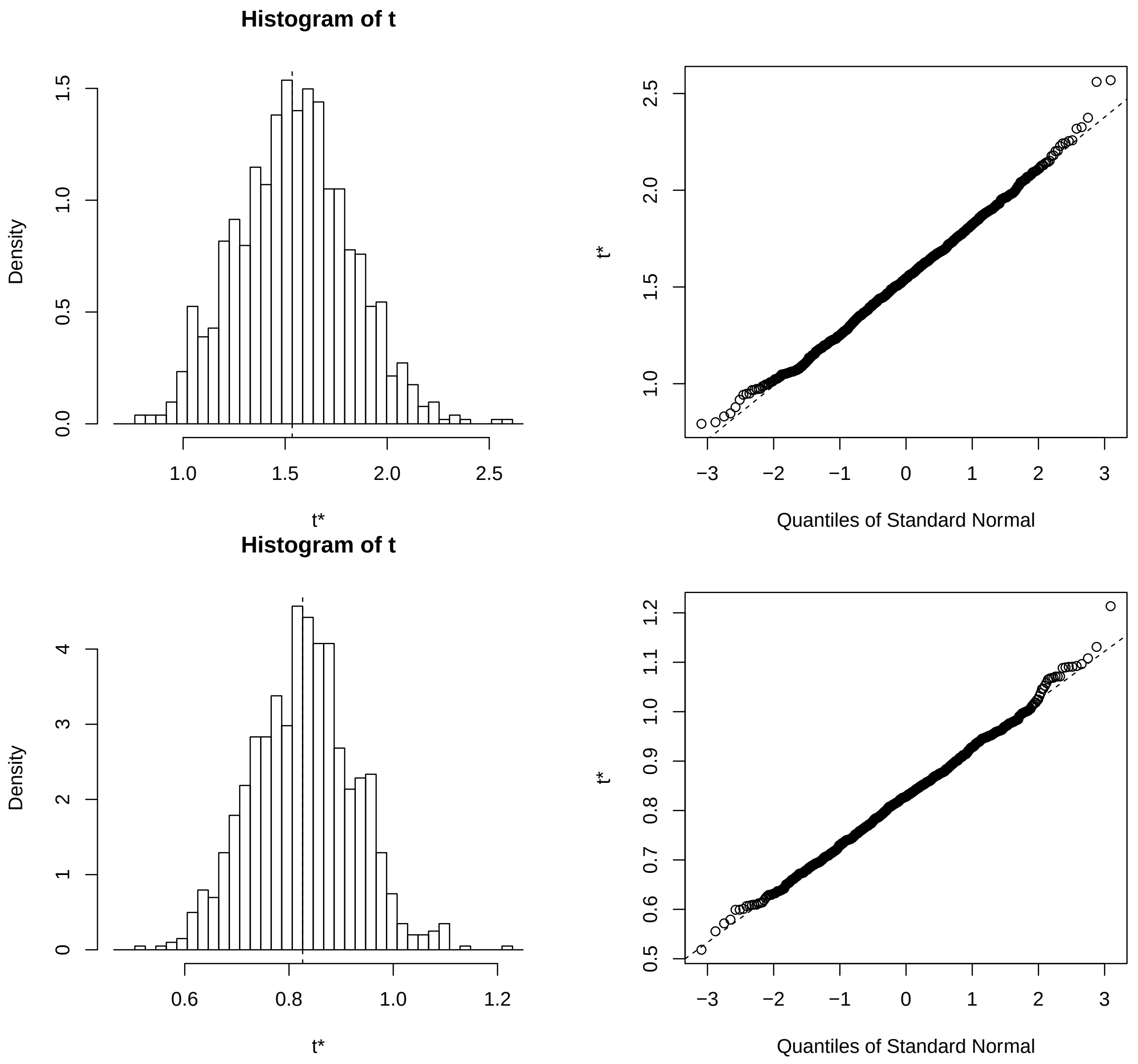

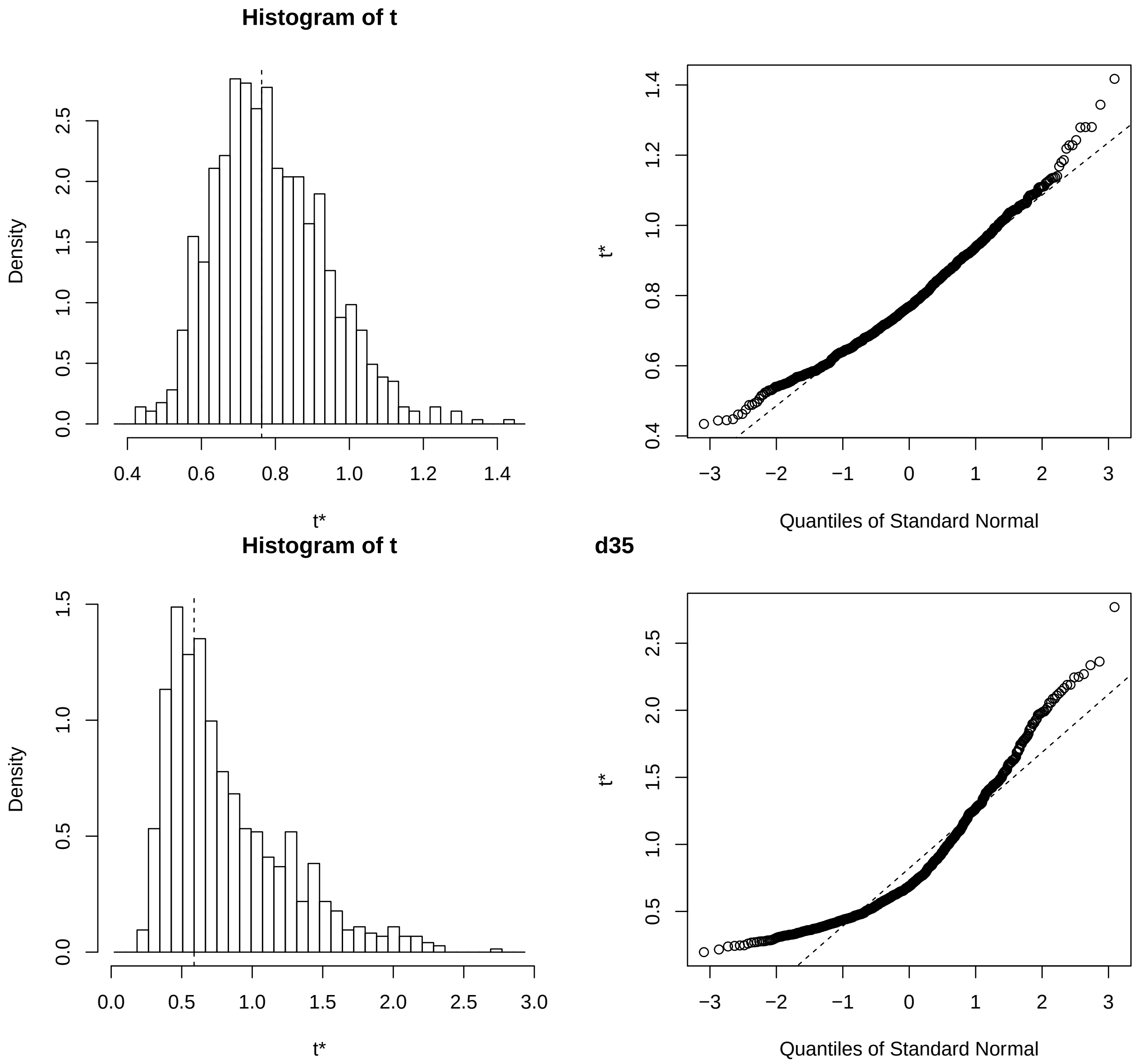

The empirical distribution of the CIs computed by resampling resembles the normal distribution for the first 3 time points, but not for the last. Nevertheless, confidence intervals assuming the normal distribution are computed for combination indices per time point:

```
vtype <- c("norm", "basic", "perc", "bca")
lapply(comb.l, boot.ci, type = vtype)
```

```
## $d14
## BOOTSTRAP CONFIDENCE INTERVAL CALCULATIONS
## Based on 1000 bootstrap replicates
##
## CALL :
## FUN(boot.out = X[[i]], type = ..1)
##
## Intervals :
## Level      Normal              Basic
## 95%   ( 0.980,  2.068 )   ( 0.970,  2.044 )
```

```
##
## Level     Percentile            BCa
## 95%   ( 1.025,  2.099 )   ( 0.974,  2.044 )
## Calculations and Intervals on Original Scale
##
## $d21
## BOOTSTRAP CONFIDENCE INTERVAL CALCULATIONS
## Based on 1000 bootstrap replicates
##
## CALL :
## FUN(boot.out = X[[i]], type = ..1)
##
## Intervals :
## Level      Normal              Basic
## 95%   ( 0.6326,  1.0174 )   ( 0.6341,  1.0185 )
##
## Level     Percentile            BCa
## 95%   ( 0.6340,  1.0185 )   ( 0.6245,  1.0005 )
## Calculations and Intervals on Original Scale
##
## $d28
## BOOTSTRAP CONFIDENCE INTERVAL CALCULATIONS
## Based on 1000 bootstrap replicates
##
## CALL :
## FUN(boot.out = X[[i]], type = ..1)
##
## Intervals :
## Level      Normal              Basic
## 95%   ( 0.4454,  1.0336 )   ( 0.4169,  0.9827 )
##
## Level     Percentile            BCa
## 95%   ( 0.5428,  1.1086 )   ( 0.5403,  1.0936 )
## Calculations and Intervals on Original Scale
##
## $d35
## BOOTSTRAP CONFIDENCE INTERVAL CALCULATIONS
## Based on 906 bootstrap replicates
##
## CALL :
## FUN(boot.out = X[[i]], type = ..1)
##
## Intervals :
## Level      Normal              Basic
## 95%   (-0.4939,  1.2001 )   (-0.7973,  0.8662 )
##
## Level     Percentile            BCa
## 95%   ( 0.3093,  1.9728 )   ( 0.2208,  1.4076 )
## Calculations and Intervals on Original Scale
## Some BCa intervals may be unstable
```

We conclude that most confidence intervals include the value 1, or values near 1, except for perhaps for one method per time point. As this is observed for different ways of computing the bootstrap confidence interval, we consider it to be robust evidence that CIs from these data do not rule out additive effect (no interaction) between the drugs.

These results can be displayed in a graph:

```
vtype <- c("norm", "basic", "perc", "bca")
# This does not work and I will no longer pursue it
cint.l <- lapply(comb.l, boot.ci, type = vtype)
vtype <- c("normal", "basic", "percent", "bca")

# Put all CInts in a matrix for plotting
cmat <- NULL
for(xi in 1:length(cint.l))
{
  for(xj in 1:length(vtype))
  {
    v.int <- cint.l[[ xi ]][ vtype[xj] ][[1]][1, ]
    cmat <- rbind(cmat, v.int[(length(v.int)-1):length(v.int)])
  }
}

rownames(cmat) <- paste( rep(vtime[2:(ntime+1)], each = length(vtype)),
                             rep(vtype, length(cint.l)) )

par(las = 2, mar = c(6, 4, 1, 1))
dcol <- var.in.colour(vtime[1:ntime], mystart = 0.55, myend = 0.85)
dcolv <- rep(dcol[[2]], each = length(vtype))

plot(1:nrow(cmat), rep(0, nrow(cmat)), col = "white", xlim = c(0.5, nrow(cmat)+0.5),
     ylim = range(cmat), xaxt = "none", xlab = "",
     ylab = "95% bootstrap confidence interval")
axis(1, at = 1:nrow(cmat), labels = rownames(cmat))
points(1:nrow(cmat), rep(combind.obs[1:ntime], each = length(vtype)), pch = 15, col = dcolv)
for(xi in 1:nrow(cmat)) segments(xi, cmat[xi, 1], xi, cmat[xi, 2], col = dcolv[xi], lwd = 2)
segments(0, 1, nrow(cmat)+0.5, 1, lty = "dotted", col = "grey", lwd = 2)
```

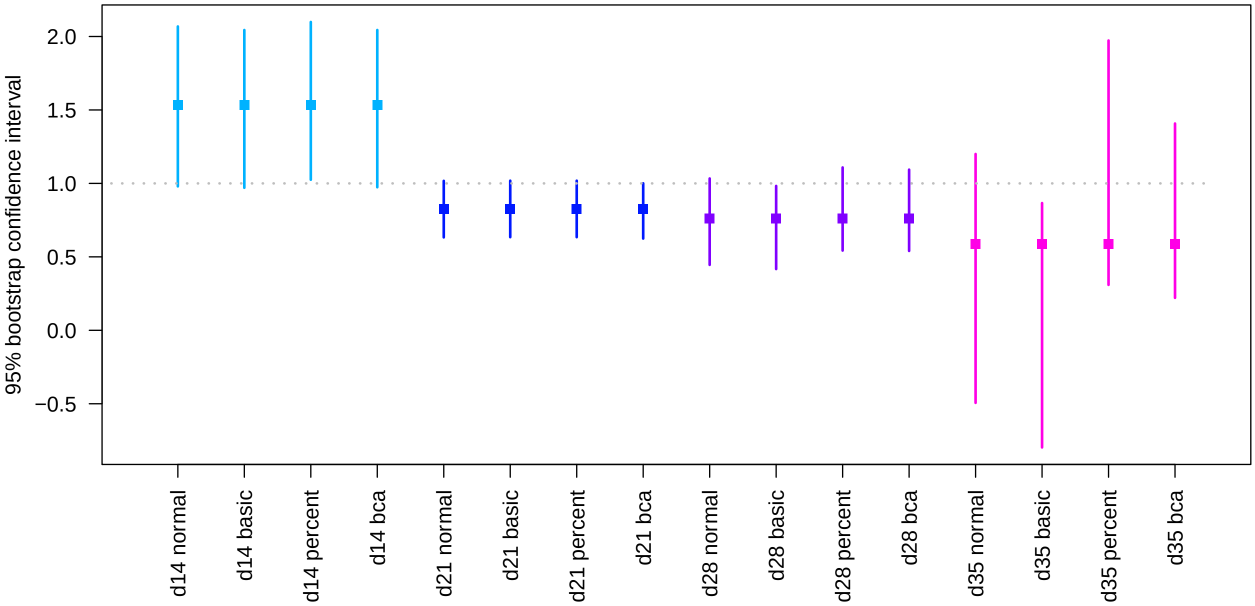

```
boot.res.d <- list(cmat, vtype, vtime, ntime, combind.obs)
```

##### Define useful group variables

For model fitting, we define two separate variables corresponding to the drug used. This enables us to model explicitly an additive as well as a non-additive (represented by an interaction, which includes but is not restricted to a synergy) effect between drugs.

```
ngroupg <- ngroupc <-  rep(0, nrow(mydata))
ngroupg[(mydata$group == levels(mydata$group)[2])|(mydata$group == levels(mydata$group)[4])] <- 1
ngroupc[(mydata$group == levels(mydata$group)[3])|(mydata$group == levels(mydata$group)[4])] <- 1
mydata$groupc <- ngroupc
mydata$groupg <- ngroupg
mydatas <- mydata # save it to use for the survival data analysis
```

##### Mixed-effects model

We need to reformat the data in order to fit the mixed-effects model. The new data matrix has one row per volume measured, and observations with `NA` are left out.

```
dnames <- paste0("d", c(7, 14, 21, 28, 35, 42, 49, 56, 63, 70, 77))
xvol <- matrix( as.matrix(mydata[, dnames]), nrow = nrow(mydata)*length(dnames), ncol = 1)[, 1]
xgroup <- factor(rep(mydata$group, length(time)), levels = levels(mydata$group))
xtime <- rep(time, each = nrow(mydata))
ngroupg <- ngroupc <-  rep(0, length(xgroup))
ngroupg[(xgroup == levels(xgroup)[2])|(xgroup == levels(xgroup)[4])] <- 1
ngroupc[(xgroup == levels(xgroup)[3])|(xgroup == levels(xgroup)[4])] <- 1
xid <- factor(rep(as.character(mydata$ID), length(time)))
lvol <- log(xvol)
lmedata <- data.frame(vol = xvol, lvol = lvol, time = xtime, group = xgroup,
                     groupg = ngroupg, groupc = ngroupc, id = xid)
lmedata <- lmedata[!is.na(lmedata$vol), ]
```

We now fit regression models with both group variables (one representing each treatment), `id` and `time`. Models assign to `id` a random effect.

```
lmefitgc <- lmer(log(vol) ~ time + groupc + groupg +  (1|id),
             REML = FALSE, na.action = na.omit, data=lmedata)
summary(lmefitgc)
```

```
## Linear mixed model fit by maximum likelihood  ['lmerMod']
## Formula: log(vol) ~ time + groupc + groupg + (1 | id)
##    Data: lmedata
##
##      AIC      BIC   logLik deviance df.resid
##    442.0    459.7   -215.0    430.0      136
##
## Scaled residuals:
##     Min      1Q  Median      3Q     Max
## -2.8330 -0.4918  0.3054  0.6525  2.7206
##
## Random effects:
##  Groups   Name        Variance Std.Dev.
##  id       (Intercept) 0.00     0.0
```

```
##  Residual             1.21     1.1
## Number of obs: 142, groups:  id, 22
##
## Fixed effects:
##              Estimate Std. Error t value
## (Intercept) 18.975803   0.205371  92.398
## time         0.119368   0.005853  20.393
## groupc      -1.085660   0.194026  -5.595
## groupg      -0.817500   0.196011  -4.171
##
## Correlation of Fixed Effects:
##        (Intr) time   groupc
## time   -0.506
## groupc -0.392 -0.252
## groupg -0.312 -0.319  0.073
## convergence code: 0
## boundary (singular) fit: see ?isSingular
```

```r
lmefitgci <- lmer(log(vol) ~ time + groupg * groupc + (1|id),
              REML = FALSE, na.action = na.omit, data=lmedata)
summary(lmefitgci)
```

```
## Linear mixed model fit by maximum likelihood  ['lmerMod']
## Formula: log(vol) ~ time + groupg * groupc + (1 | id)
##    Data: lmedata
##
##      AIC      BIC   logLik deviance df.resid
##    444.0    464.7   -215.0    430.0      135
##
## Scaled residuals:
##     Min      1Q  Median      3Q     Max
## -2.8446 -0.5065  0.3178  0.6507  2.7414
##
## Random effects:
##  Groups   Name        Variance Std.Dev.
##  id       (Intercept) 0.000    0.0
##  Residual             1.209    1.1
## Number of obs: 142, groups:  id, 22
##
## Fixed effects:
##               Estimate Std. Error t value
## (Intercept)   19.00404    0.24230  78.433
## time           0.11930    0.00586  20.358
## groupg        -0.86584    0.29481  -2.937
## groupc        -1.13118    0.28398  -3.983
## groupg:groupc  0.08303    0.37827   0.219
##
## Correlation of Fixed Effects:
##              (Intr) time   groupg groupc
## time         -0.456
## groupg       -0.572 -0.173
## groupc       -0.615 -0.134  0.579
## groupg:grpc   0.531 -0.051 -0.747 -0.730
## convergence code: 0
## boundary (singular) fit: see ?isSingular
```

```r
anova(lmefitgc, lmefitgci)
```

```
## Data: lmedata
## Models:
## lmefitgc: log(vol) ~ time + groupc + groupg + (1 | id)
## lmefitgci: log(vol) ~ time + groupg * groupc + (1 | id)
##           npar    AIC    BIC  logLik deviance  Chisq Df Pr(>Chisq)
## lmefitgc     6 442.01 459.75 -215.01   430.01
## lmefitgci    7 443.96 464.65 -214.98   429.96 0.0482  1     0.8263
```

The ANOVA test leads us to conclude that the model fit including the interaction effect is not better than the model including only the main (additive) effects of the individual drugs.

##### Survival model

We also study the impact of treatments additively, as well as including their interaction, on the overall survival.

```r
mydata <- mydatas
msurv <- Surv(mydata$endpoint)

mcox <- coxph(msurv ~ groupc + groupg + cluster(ID), x = TRUE, model = TRUE, data = mydata)
summary(mcox)
```

```
## Call:
## coxph(formula = msurv ~ groupc + groupg, data = mydata, model = TRUE,
##     x = TRUE, cluster = ID)
##
##   n= 22, number of events= 22
##
##            coef exp(coef) se(coef) robust se      z Pr(>|z|)
## groupc -1.77607   0.16930  0.56646   0.43350 -4.097 4.18e-05 ***
## groupg -2.50327   0.08182  0.72351   0.74487 -3.361 0.000777 ***
## ---
## Signif. codes:  0 '***' 0.001 '**' 0.01 '*' 0.05 '.' 0.1 ' ' 1
##
##        exp(coef) exp(-coef) lower .95 upper .95
## groupc   0.16930      5.907   0.07239    0.3960
## groupg   0.08182     12.222   0.01900    0.3523
##
## Concordance= 0.782  (se = 0.051 )
## Likelihood ratio test= 21.07  on 2 df,   p=3e-05
## Wald test            = 20.52  on 2 df,   p=4e-05
## Score (logrank) test = 19.44  on 2 df,   p=6e-05,   Robust = 15.28  p=5e-04
##
##   (Note: the likelihood ratio and score tests assume independence of
##      observations within a cluster, the Wald and robust score tests do not).
```

```r
mcoxi <- coxph(msurv ~ groupc * groupg + cluster(ID), x = TRUE, model = TRUE, data = mydata)
summary(mcoxi)
```

```
## Call:
## coxph(formula = msurv ~ groupc + groupg + groupc:groupg, data = mydata,
##     model = TRUE, x = TRUE, cluster = ID)
##
```

```
##   n= 22, number of events= 22
##
##                    coef exp(coef) se(coef) robust se      z Pr(>|z|)
## groupc        -1.79554   0.16604  0.75186   0.57725 -3.110  0.00187 **
## groupg        -2.52415   0.08013  0.89714   1.04508 -2.415  0.01572 *
## groupc:groupg  0.04436   1.04536  1.12242   1.03486  0.043  0.96581
## ---
## Signif. codes:  0 '***' 0.001 '**' 0.01 '*' 0.05 '.' 0.1 ' ' 1
##
##               exp(coef) exp(-coef) lower .95 upper .95
## groupc          0.16604     6.0227   0.05356    0.5147
## groupg          0.08013    12.4803   0.01033    0.6214
## groupc:groupg   1.04536     0.9566   0.13753    7.9459
##
## Concordance= 0.782  (se = 0.051 )
## Likelihood ratio test= 21.07  on 3 df,   p=1e-04
## Wald test            = 20.72  on 3 df,   p=1e-04
## Score (logrank) test = 22.4  on 3 df,   p=5e-05,   Robust = 15.51  p=0.001
##
##   (Note: the likelihood ratio and score tests assume independence of
##      observations within a cluster, the Wald and robust score tests do not).
```

So, in terms of survival, the two drugs yield statistically different progressions from that of mice in the control group, independently and additively. However, an added interaction effect between drugs to the model, which could capture effects that cannot be explained additively (including synergy, but not exclusively so), is not statistically significant. This leads us to conclude that, on the basis of the current data, there is no evidence that these two drugs have a synergistic effect on progression.

Note that we fit Cox proportional-hazards models with robust variance estimates, via the option `cluster`.

Kaplan-Meier survival curves for all animals, as well as per group, are displayed below.

```
gcol <- var.in.colour(mydata$group)
par(mfrow=c(1, 2))
plot(msurv, main =  "Kaplan-Meier", ylab = "estimated survival function", col = "blue",
     xlab = "days")
plot(survfit(msurv ~ mydata$group), main =  "Kaplan-Meier per group", ylab = "estimated survival functi
     xlab = "days")
legend("bottomleft", legend = gcol[[3]], lty = "solid", col = gcol[[2]], cex = .6)
```

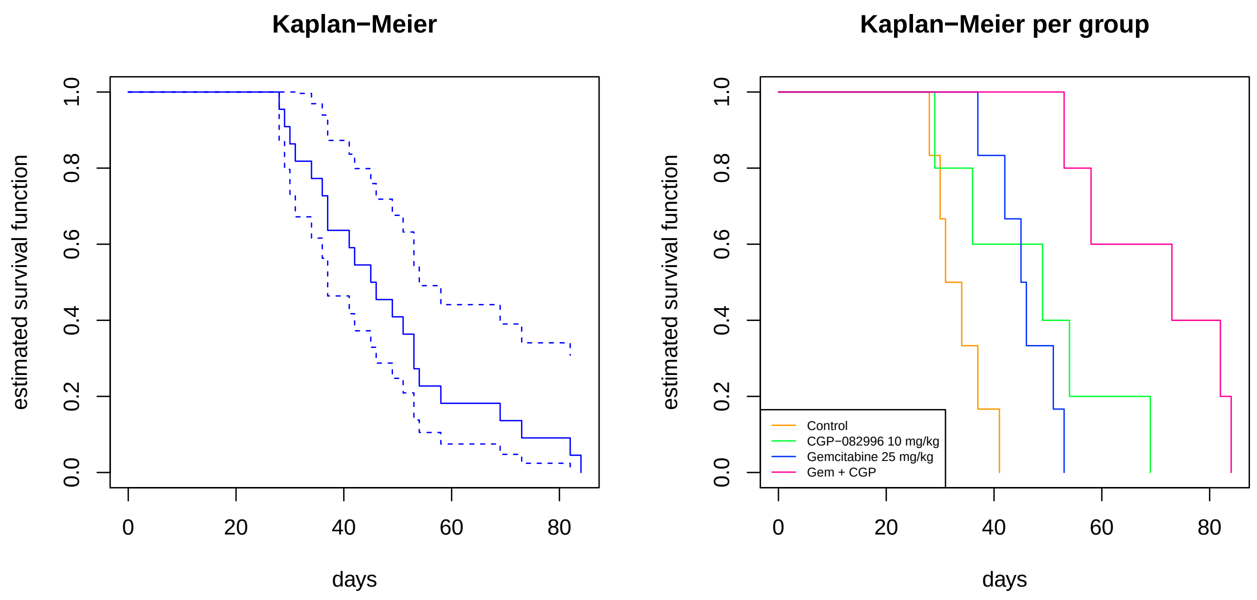

#### Figure 5C

##### Read in and explore data

```
mydata <- read.delim("fig5c.txt")
time <- c(16, 23, 31, 37, 44, 51, 58, 65, 72, 79, 86)
mydata0 <- mydata # with original column names, to be used in the survival data analysis
colnames(mydata)[3:(ncol(mydata)-1)] <- paste0("d", time)
mydata$group <- factor(mydata$group, levels = c("Control", "AZ628 10 mg/kg",
                                                "Gemcitabine 50 mg/kg", "AZ628 + Gemcitabine"))
```

The data read in contains information on 30 animals measured by 14 variables. Note that there are 2 animals with no tumor growth, which we now leave out of analyses.

```
mydata <- mydata[ rowSums(is.na(mydata)) < 12, ]
```

Some variables have many `NA` entries:

```
colSums(is.na(mydata))
```

```
##       ID    group      d16      d23      d31      d37      d44      d51
##        0        0        0        0        0        2        9       12
##      d58      d65      d72      d79      d86 endpoint
##       15       18       18       21       25        0
```

Visualizing the data per animal, within groups:

```
gcol <- var.in.colour(mydata$group, mystart = 0.1, myend = 0.9)
par(mfrow=c(1, 4), bg = "grey90")
for(xg in 1:nlevels(mydata$group))
{
    datag <- mydata[ mydata$group == levels(mydata$group)[ xg ], ]
    dplot <- datag[, 3:(ncol(datag)-1)]
    plot(1, 1, col = "white", xlim = range(time), ylim = range(dplot, na.rm = TRUE),
         xlab = "time (days)", ylab = "tumor volume", main = levels(mydata$group)[ xg ])
```

```
    for(xr in 1:nrow(datag)) lines(time, dplot[xr, ], col = gcol[[2]][ gcol[[3]] == levels(mydata$group)
    }
```

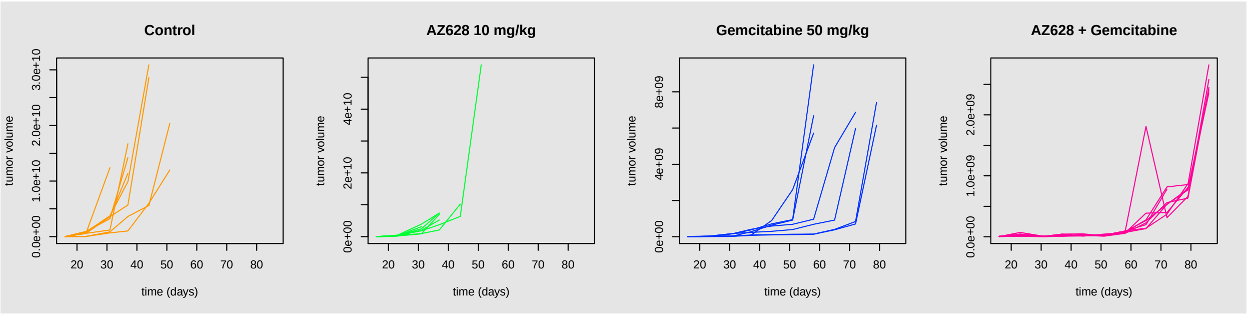

For these data, the initial value can be taken as $T = 16$. We divide the volume values by the one at $T = 16$ per mouse and log-transform the data. Then the tumor volume evolution for animals coloured by group is:

```
gcol <- var.in.colour(mydata$group, mystart = 0.1, myend = 0.9)
par(bg = "grey90")
data.all <- mydata[, 3:(ncol(mydata)-1)]
data.all <- data.all/matrix(data.all[, 1], nrow = nrow(data.all), ncol = ncol(data.all))
myylim <- range(log10(data.all), na.rm=TRUE)
plot(1, 1, col = "white", xlim = range(time), ylim = myylim,  # ylim = range(log10(dplot),na.rm = TRUE)
         xlab = "time (days)", ylab = "tumor volume (log10 %)")
for(xg in 1:nlevels(mydata$group))
{
    datag <- data.all[ mydata$group == levels(mydata$group)[ xg ], ]
    dplot <- datag
    for(xr in 1:nrow(datag)) lines(time, log10(dplot[xr, ]), col = gcol[[2]][ gcol[[3]] == levels(mydat
    }
legend("bottomright", legend = gcol[[3]], lty = "solid", col = gcol[[2]], cex = .8)
```

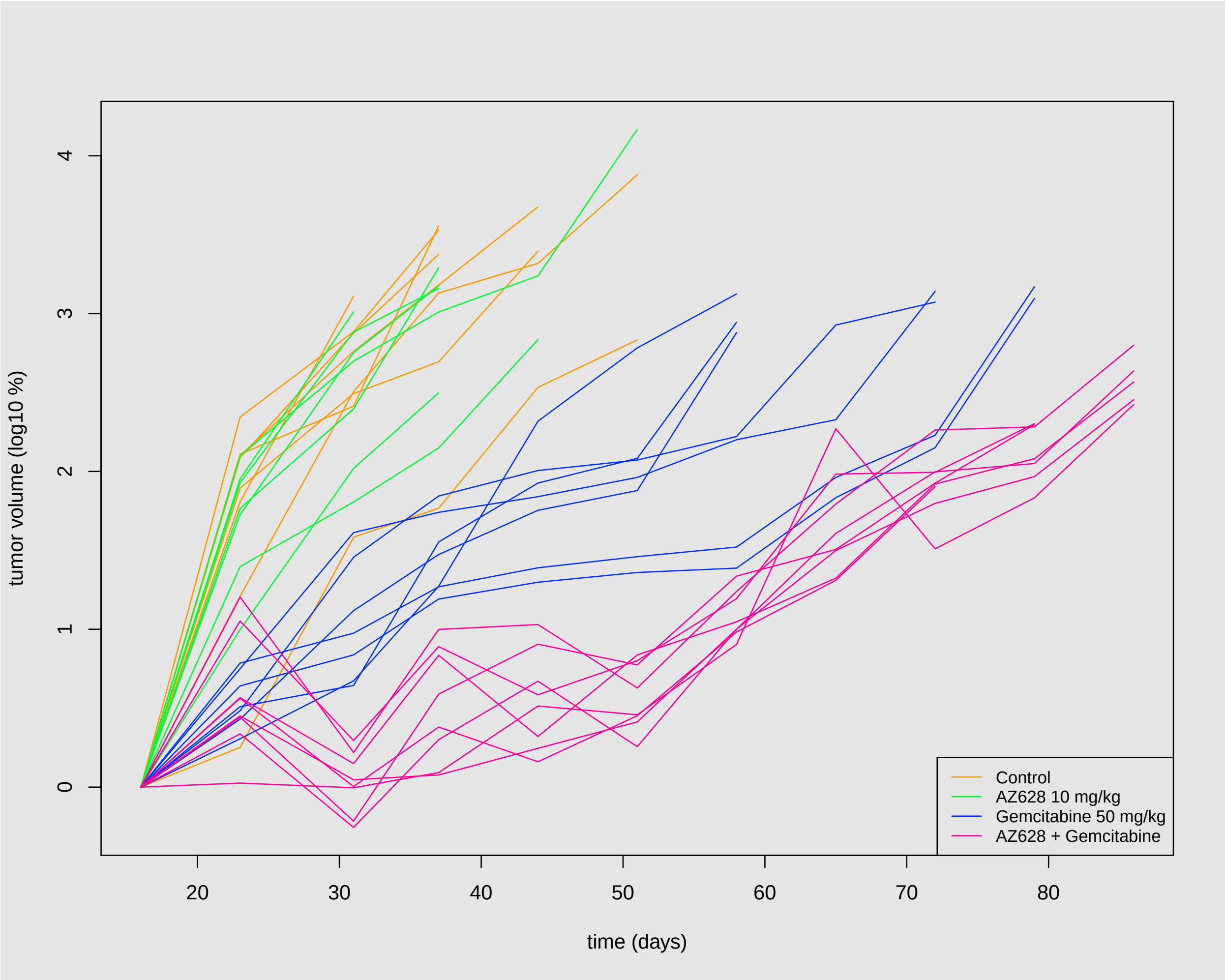

```
mydatac <- mydata
```

##### Computing the Combination Index

The combination index was computed in the original paper as follows:

- computed tumor volume average per time point and group
- divide each average by that for the initial time point
- divide each tumor volume percentage by that for the control group (group of mice which did not receive any of the drugs)

We will recompute the CIs per time point using the data for Figure 5C. Subsequently, we will construct confidence intervals for the CIs using the stratified bootstrap, which performs resampling taking the groups into account.

The Combination Indices for the original data, with no data transformation, per time point, are:

```
combind.obs <- combind(mydata, xi = 1:nrow(mydata))

round(combind.obs, 3)
```

```
##      d23     d31     d37     d44     d51     d58     d65     d72
##    1.353   0.082   0.112   0.054   0.030     NaN     NaN     NaN
```

```
##       d79       d86  endpoint
##       NaN       NaN     3.608
```

In this case, only the first 5 time points (discarding the initial time point, which serves as a reference) have valid results for the CIs. Indeed, for later time points all individuals in the control group were no longer available.

We now build confidence intervals for Combination Indices using the stratified bootstrap with 1000 resamples. Only CIs with valid results are used, so this is restricted to the first 5 time points. Here we use the function `boot` from the package of the same name to do this. The stratified bootstrap performs resampling within groups defined by the treatment, thereby ensuring that groups are always well represented in each resample.

```
vtime <- paste0("d", time)
ntime <- 5 # number of time points used, after the reference one
comb.l <- vector("list", ntime)
names(comb.l) <- vtime[2:(ntime + 1)]
for(xi in 2:(ntime+1))
{
  mydatat <- cbind(mydata[, 1:3],  mydata[, vtime[xi] ])
  names(mydatat)[ncol(mydatat)] <- vtime[xi]
  comb.l[[ vtime[xi] ]] <- boot(mydatat, statistic = combind.boot, R = 1000, stype = "i", strata = myda
}
```

Per time point, the empirical distribution of the indices is:

```
for(xi in 1:ntime) plot(comb.l[[ xi ]]); title( paste(vtime[xi + 1]))
```

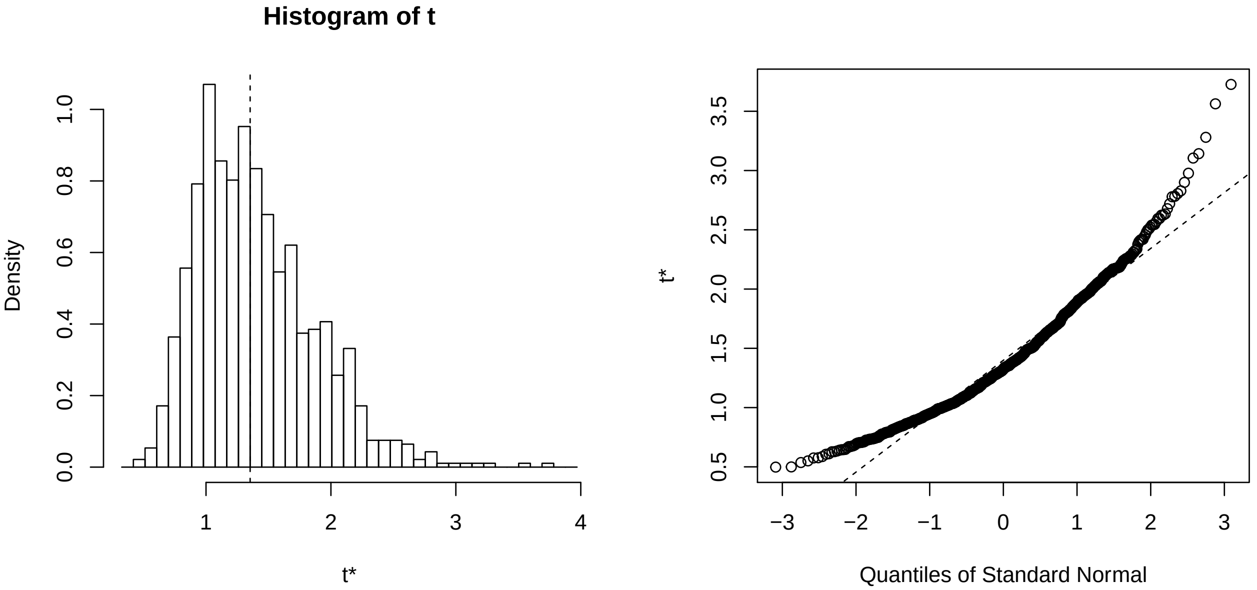

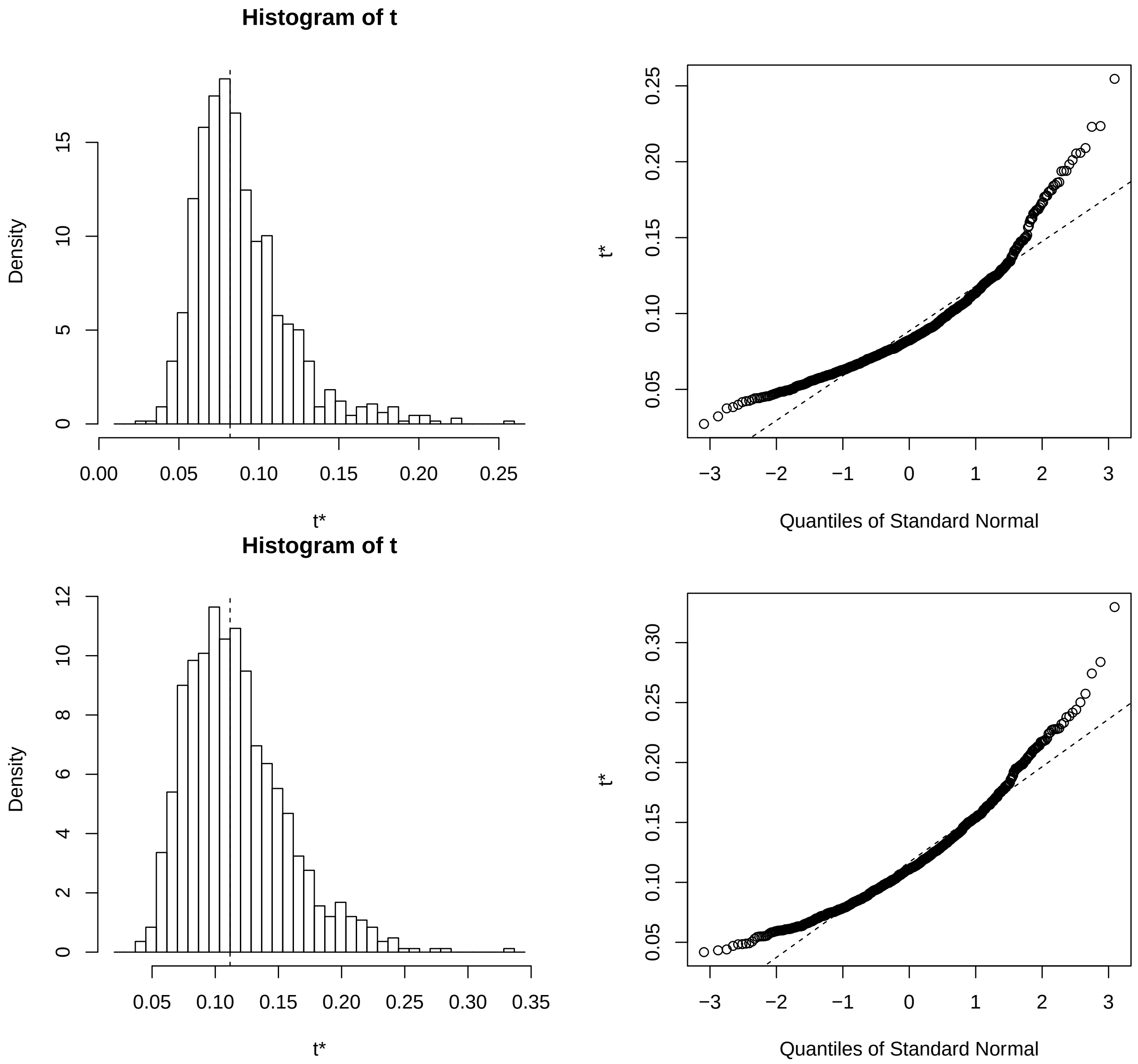
Histogram of t
Density
t*
Quantiles of Standard Normal
Histogram of t
Density
t*
t*
Quantiles of Standard Normal

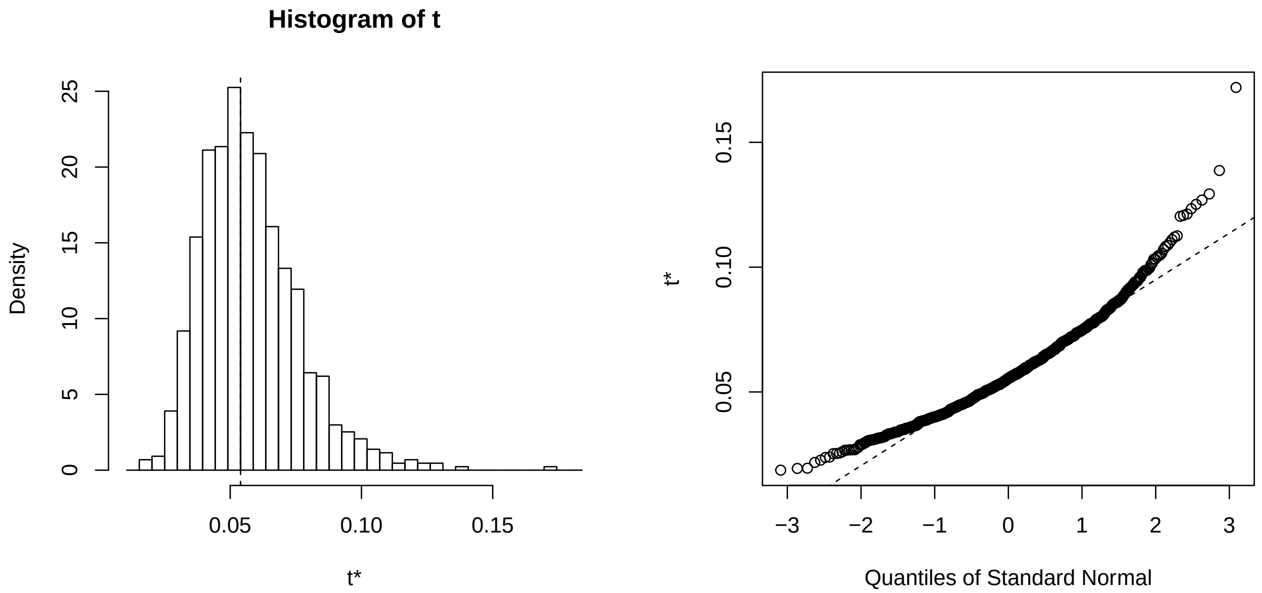

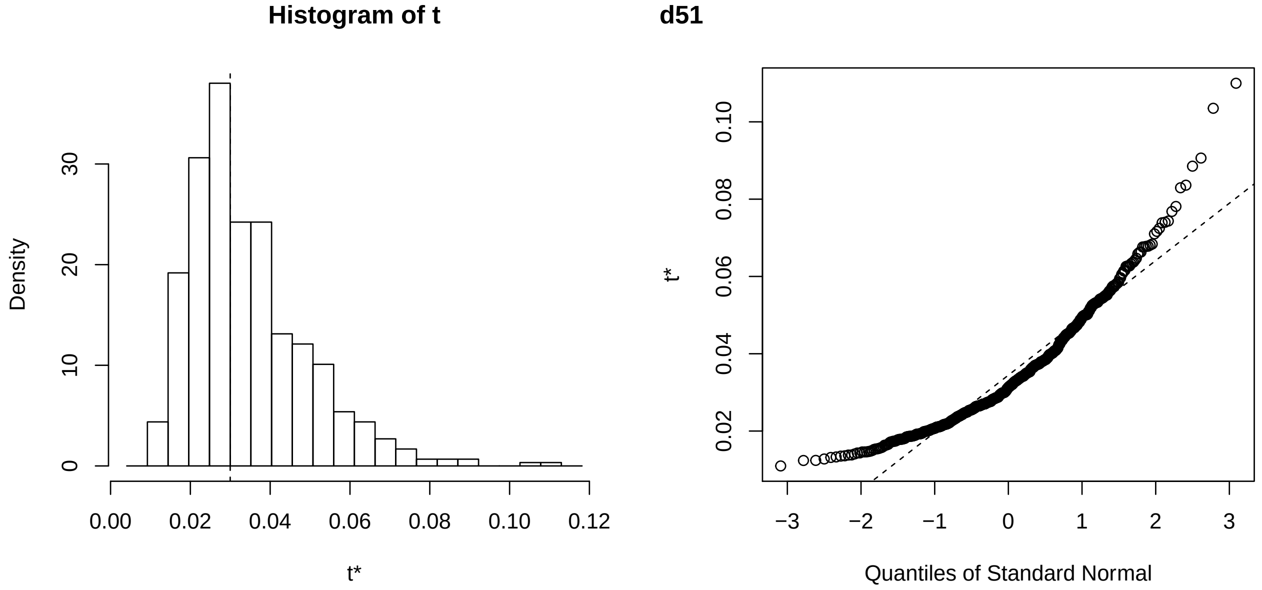

The empirical distribution of the CIs computed by resampling does not resemble the normal distribution for any of the time points, in this case, being left-skewed. Nevertheless, confidence intervals were computed, including ones assuming the normal distribution, for combination indices per time point:

```r
vtype <- c("norm", "basic", "perc", "bca")
lapply(comb.l, boot.ci, type = vtype)
```

```
## $d23
## BOOTSTRAP CONFIDENCE INTERVAL CALCULATIONS
## Based on 1000 bootstrap replicates
##
## CALL :
## FUN(boot.out = X[[i]], type = ..1)
##
## Intervals :
## Level      Normal              Basic
## 95%   ( 0.383,  2.232 )   ( 0.206,  2.003 )
```

```
##
## Level     Percentile            BCa
## 95%   ( 0.704,  2.500 )   ( 0.772,  2.783 )
## Calculations and Intervals on Original Scale
## Some BCa intervals may be unstable
##
## $d31
## BOOTSTRAP CONFIDENCE INTERVAL CALCULATIONS
## Based on 1000 bootstrap replicates
##
## CALL :
## FUN(boot.out = X[[i]], type = ..1)
##
## Intervals :
## Level      Normal              Basic
## 95%   ( 0.0176,  0.1333 )   (-0.0059,  0.1159 )
##
## Level     Percentile            BCa
## 95%   ( 0.0481,  0.1699 )   ( 0.0455,  0.1578 )
## Calculations and Intervals on Original Scale
##
## $d37
## BOOTSTRAP CONFIDENCE INTERVAL CALCULATIONS
## Based on 1000 bootstrap replicates
##
## CALL :
## FUN(boot.out = X[[i]], type = ..1)
##
## Intervals :
## Level      Normal              Basic
## 95%   ( 0.0286,  0.1845 )   ( 0.0093,  0.1637 )
##
## Level     Percentile            BCa
## 95%   ( 0.0597,  0.2141 )   ( 0.0600,  0.2178 )
## Calculations and Intervals on Original Scale
##
## $d44
## BOOTSTRAP CONFIDENCE INTERVAL CALCULATIONS
## Based on 904 bootstrap replicates
##
## CALL :
## FUN(boot.out = X[[i]], type = ..1)
##
## Intervals :
## Level      Normal              Basic
## 95%   ( 0.0136,  0.0867 )   ( 0.0057,  0.0785 )
##
## Level     Percentile            BCa
## 95%   ( 0.0294,  0.1022 )   ( 0.0271,  0.0985 )
## Calculations and Intervals on Original Scale
##
## $d51
## BOOTSTRAP CONFIDENCE INTERVAL CALCULATIONS
## Based on 573 bootstrap replicates
```

```
##
## CALL :
## FUN(boot.out = X[[i]], type = ..1)
##
## Intervals :
## Level      Normal              Basic
## 95%   (-0.0036,  0.0546 )   (-0.0101,  0.0454 )
##
## Level     Percentile            BCa
## 95%   ( 0.0146,  0.0701 )   ( 0.0132,  0.0630 )
## Calculations and Intervals on Original Scale
## Some BCa intervals may be unstable
```

We conclude that most confidence intervals include the value 1, or values near enough one. As this is observed for different ways of computing the bootstrap confidence interval, we consider it to be robust evidence that CIs from these data do not rule out additive effect (no interaction) between the drugs.

We conclude that confidence intervals do not include value 1, or values near 1, except for the second time point (first CI). As this is observed for different ways of computing the bootstrap confidence interval, we consider it to be robust evidence that CIs from these data rule out additive effect (no interaction) between the drugs.

These results can be displayed in a graph:

```
vtype <- c("norm", "basic", "perc", "bca")
cint.l <- lapply(comb.l, boot.ci, type = vtype)
vtype <- c("normal", "basic", "percent", "bca")

# Put all CInts in a matrix for plotting
cmat <- NULL
for(xi in 1:length(cint.l))
{
  for(xj in 1:length(vtype))
  {
    v.int <- cint.l[[ xi ]][ vtype[xj] ][[1]][1, ]
    cmat <- rbind(cmat, v.int[(length(v.int)-1):length(v.int)])
  }
}

rownames(cmat) <- paste( rep(vtime[2:(ntime+1)], each = length(vtype)),
                             rep(vtype, length(cint.l)) )

par(las = 2, mar = c(6, 4, 1, 1))
dcol <- var.in.colour(vtime[1:ntime], mystart = 0.55, myend = 0.85)
dcolv <- rep(dcol[[2]], each = length(vtype))

plot(1:nrow(cmat), rep(0, nrow(cmat)), col = "white", xlim = c(0.5, nrow(cmat)+0.5),
     ylim = range(cmat), xaxt = "none", xlab = "",
     ylab = "95% bootstrap confidence interval")
axis(1, at = 1:nrow(cmat), labels = rownames(cmat))
points(1:nrow(cmat), rep(combind.obs[1:ntime], each = length(vtype)), pch = 15, col = dcolv)
for(xi in 1:nrow(cmat)) segments(xi, cmat[xi, 1], xi, cmat[xi, 2], col = dcolv[xi], lwd = 2)
segments(0, 1, nrow(cmat)+0.5, 1, lty = "dotted", col = "grey", lwd = 2)
```

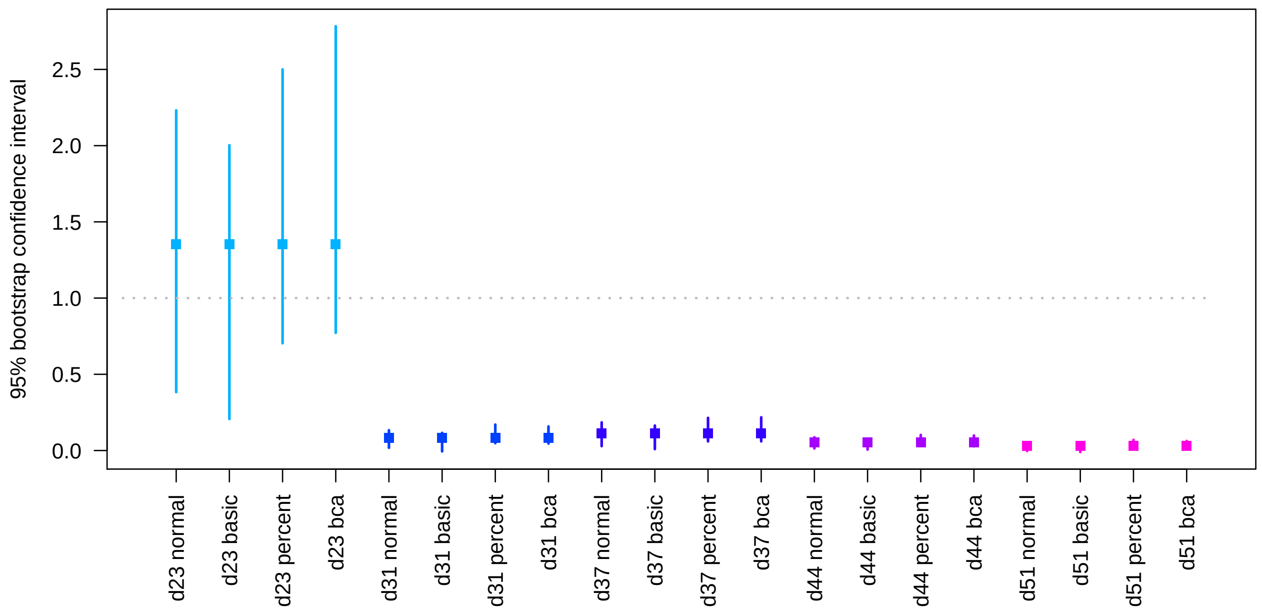

```
boot.res.c <- list(cmat, vtype, vtime, ntime, combind.obs)
```

##### Define useful group variables

For model fitting, we define two separate variables corresponding to the drug used. This enables us to model explicitly an additive as well as a non-additive (represented by an interaction, which includes but is not restricted to a synergy) effect between drugs.

```
ngroupg <- ngroupc <-  rep(0, nrow(mydata))
ngroupg[(mydata$group == levels(mydata$group)[2])|(mydata$group == levels(mydata$group)[4])] <- 1
ngroupc[(mydata$group == levels(mydata$group)[3])|(mydata$group == levels(mydata$group)[4])] <- 1
mydata$groupc <- ngroupc
mydata$groupg <- ngroupg
mydatas <- mydata # save it to use for the survival data analysis
```

##### Mixed-effects model

We need to reformat the data in order to fit the mixed-effects model. The new data matrix has one row per volume measured, and observations with `NA` are left out.

```
#leave out the endpoint for this analysis
dnames <- colnames(mydata)[3:(2+length(time))] # paste0("X", time)
xvol <- matrix( as.matrix(mydata[, dnames]), nrow = nrow(mydata)*length(dnames), ncol = 1)[, 1]
xgroup <- factor(rep(mydata$group, length(time)), levels = levels(mydata$group))
xtime <- rep(time, each = nrow(mydata))
ngroupg <- ngroupc <-  rep(0, length(xgroup))
ngroupg[(xgroup == levels(xgroup)[2])|(xgroup == levels(xgroup)[4])] <- 1
ngroupc[(xgroup == levels(xgroup)[3])|(xgroup == levels(xgroup)[4])] <- 1
xid <- factor(rep(as.character(mydata$ID), length(time)))
lvol <- log(xvol)
lmedata <- data.frame(vol = xvol, lvol = lvol, time = xtime, group = xgroup,
                      groupg = ngroupg, groupc = ngroupc, id = xid)
lmedata <- lmedata[!is.na(lmedata$vol), ]
```

We now fit regression models with both group variables (one representing each treatment), `id` and `time`. Models assign to `id` a random effect.

```
lmefitgc <- lmer(log(vol) ~ time + groupc + groupg +  (1|id),
              REML = FALSE, na.action = na.omit, data=lmedata)
summary(lmefitgc)
```

```
## Linear mixed model fit by maximum likelihood  ['lmerMod']
## Formula: log(vol) ~ time + groupc + groupg + (1 | id)
##    Data: lmedata
##
##      AIC      BIC   logLik deviance df.resid
##    753.4    773.5   -370.7    741.4      204
##
## Scaled residuals:
##     Min      1Q  Median      3Q     Max
## -3.1808 -0.5214  0.0151  0.6959  1.9769
##
## Random effects:
##  Groups   Name        Variance Std.Dev.
##  id       (Intercept) 0.06971  0.264
##  Residual             1.93673  1.392
## Number of obs: 210, groups:  id, 30
##
## Fixed effects:
##              Estimate Std. Error t value
## (Intercept) 17.746640   0.256696   69.14
## time         0.104803   0.005498   19.06
## groupc      -3.465393   0.248192  -13.96
## groupg      -1.307208   0.220794   -5.92
##
## Correlation of Fixed Effects:
##        (Intr) time   groupc
## time   -0.582
## groupc -0.266 -0.379
## groupg -0.327 -0.098 -0.072
```

```
lmefitgci <- lmer(log(vol) ~ time + groupg * groupc + (1|id),
              REML = FALSE, na.action = na.omit, data=lmedata)
summary(lmefitgci)
```

```
## Linear mixed model fit by maximum likelihood  ['lmerMod']
## Formula: log(vol) ~ time + groupg * groupc + (1 | id)
##    Data: lmedata
##
##      AIC      BIC   logLik deviance df.resid
##    741.5    764.9   -363.7    727.5      203
##
## Scaled residuals:
##     Min      1Q  Median      3Q     Max
## -3.0210 -0.5255 -0.0094  0.7298  2.1740
##
## Random effects:
##  Groups   Name        Variance Std.Dev.
##  id       (Intercept) 0.000    0.000
```

```
##  Residual             1.871    1.368
## Number of obs: 210, groups:  id, 30
##
## Fixed effects:
##                Estimate Std. Error t value
## (Intercept)   17.240296   0.276167  62.427
## time           0.106047   0.005408  19.610
## groupg        -0.265029   0.336108  -0.789
## groupc        -2.708824   0.295530  -9.166
## groupg:groupc -1.593773   0.410807  -3.880
##
## Correlation of Fixed Effects:
##             (Intr) time   groupg groupc
## time        -0.581
## groupg      -0.557  0.021
## groupc      -0.480 -0.241  0.504
## groupg:grpc  0.507 -0.105 -0.820 -0.652
## convergence code: 0
## boundary (singular) fit: see ?isSingular
```

```r
anova(lmefitgc, lmefitgci)
```

```
## Data: lmedata
## Models:
## lmefitgc: log(vol) ~ time + groupc + groupg + (1 | id)
## lmefitgci: log(vol) ~ time + groupg * groupc + (1 | id)
##           npar    AIC    BIC  logLik deviance  Chisq Df Pr(>Chisq)
## lmefitgc     6 753.41 773.49 -370.70   741.41
## lmefitgci    7 741.49 764.92 -363.74   727.49 13.919  1  0.0001909 ***
## ---
## Signif. codes:  0 '***' 0.001 '**' 0.01 '*' 0.05 '.' 0.1 ' ' 1
```

The ANOVA test leads us to conclude that the model fit including the interaction effect is better than the model including only the main (additive) effects of the individual drugs.

##### Survival model

We also study the impact of treatments additively, as well as including their interaction, on the overall survival.

```r
mydata <- mydatas
msurv <- Surv(mydata$endpoint)

mcox <- coxph(msurv ~ groupc + groupg + cluster(ID), x = TRUE, model = TRUE, data = mydata)
```

```
## Warning in fitter(X, Y, istrat, offset, init, control, weights = weights, :
## Loglik converged before variable 1 ; coefficient may be infinite.
```

```r
summary(mcox)
```

```
## Call:
## coxph(formula = msurv ~ groupc + groupg, data = mydata, model = TRUE,
##     x = TRUE, cluster = ID)
##
##   n= 30, number of events= 30
##
```

```
##              coef  exp(coef)   se(coef)  robust se       z Pr(>|z|)
## groupc -2.156e+01  4.334e-10  7.244e+03  3.230e-01 -66.741   <2e-16 ***
## groupg -5.413e-01  5.820e-01  4.529e-01  3.896e-01  -1.389    0.165
## ---
## Signif. codes:  0 '***' 0.001 '**' 0.01 '*' 0.05 '.' 0.1 ' ' 1
##
##        exp(coef) exp(-coef) lower .95 upper .95
## groupc 4.334e-10  2.308e+09 2.301e-10 8.162e-10
## groupg 5.820e-01  1.718e+00 2.712e-01 1.249e+00
##
## Concordance= 0.809  (se = 0.037 )
## Likelihood ratio test= 39.18  on 2 df,   p=3e-09
## Wald test            = 5049  on 2 df,   p=<2e-16
## Score (logrank) test = 35.86  on 2 df,   p=2e-08,   Robust = 27.17  p=1e-06
##
##   (Note: the likelihood ratio and score tests assume independence of
##      observations within a cluster, the Wald and robust score tests do not).
```

```
mcoxi <- coxph(msurv ~ groupc * groupg + cluster(ID), x = TRUE, model = TRUE, data = mydata)
```

```
## Warning in fitter(X, Y, istrat, offset, init, control, weights = weights, :
## Loglik converged before variable 1 ; coefficient may be infinite.
```

```
summary(mcoxi)
```

```
## Call:
## coxph(formula = msurv ~ groupc + groupg + groupc:groupg, data = mydata,
##     model = TRUE, x = TRUE, cluster = ID)
##
##   n= 30, number of events= 30
##
##                    coef  exp(coef)   se(coef)  robust se       z Pr(>|z|)
## groupc        -2.087e+01  8.649e-10  7.044e+03  4.224e-01 -49.409  < 2e-16 ***
## groupg         4.611e-01  1.586e+00  5.394e-01  5.102e-01   0.904  0.36612
## groupc:groupg -2.478e+00  8.394e-02  9.025e-01  7.830e-01  -3.164  0.00155 **
## ---
## Signif. codes:  0 '***' 0.001 '**' 0.01 '*' 0.05 '.' 0.1 ' ' 1
##
##               exp(coef) exp(-coef) lower .95 upper .95
## groupc        8.649e-10  1.156e+09 3.779e-10 1.979e-09
## groupg        1.586e+00  6.306e-01 5.834e-01 4.311e+00
## groupc:groupg 8.394e-02  1.191e+01 1.809e-02 3.895e-01
##
## Concordance= 0.847  (se = 0.028 )
## Likelihood ratio test= 47.12  on 3 df,   p=3e-10
## Wald test            = 3372  on 3 df,   p=<2e-16
## Score (logrank) test = 39.65  on 3 df,   p=1e-08,   Robust = 28.09  p=3e-06
##
##   (Note: the likelihood ratio and score tests assume independence of
##      observations within a cluster, the Wald and robust score tests do not).
```

So, in terms of survival, the two drugs yield statistically different progressions from that of mice in the control group, independently and additively. In addition, the interaction effect between the two drugs, which captures effects that cannot be explained additively (including synergy, but not exclusively so), is statistically significant. This leads us to conclude that, on the basis of the current data, there is evidence that these two drugs have a synergistic effect on progression.

Note that we fit Cox proportional-hazards models with robust variance estimates, via the option `cluster`.

Kaplan-Meier survival curves for all animals, as well as per group, are displayed below.

```
gcol <- var.in.colour(mydata$group)
par(mfrow=c(1, 2))
plot(msurv, main =  "Kaplan-Meier", ylab = "estimated survival function",
     col = "blue", xlab = "days")
plot(survfit(msurv ~ mydata$group), main =  "Kaplan-Meier per group",
     ylab = "estimated survival function", col = gcol[[2]],
     xlab = "days")
legend("bottomleft", legend = gcol[[3]], lty = "solid", col = gcol[[2]], cex = .6)
```

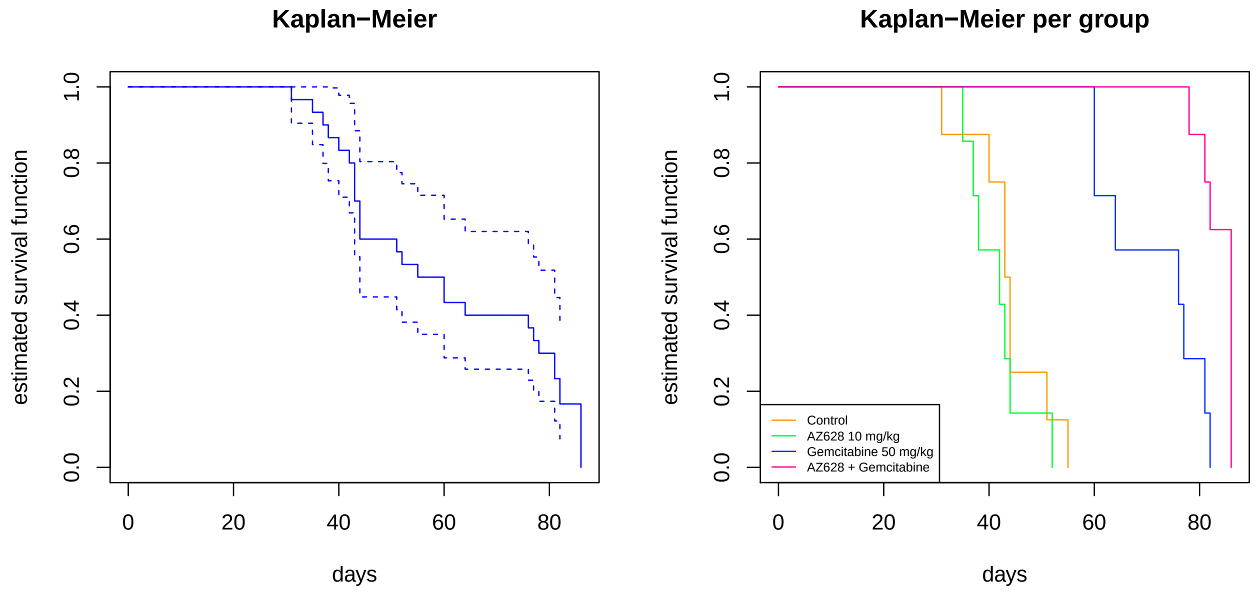

#### Package versions

For completeness, packages used and their versions are listed below:

```
sessionInfo()
```

```
## R version 3.6.3 (2020-02-29)
## Platform: x86_64-pc-linux-gnu (64-bit)
## Running under: Ubuntu 18.04.5 LTS
##
## Matrix products: default
## BLAS:   /usr/lib/x86_64-linux-gnu/blas/libblas.so.3.7.1
## LAPACK: /usr/lib/x86_64-linux-gnu/lapack/liblapack.so.3.7.1
##
## locale:
##  [1] LC_CTYPE=en_US.UTF-8       LC_NUMERIC=C
##  [3] LC_TIME=nl_NL.UTF-8        LC_COLLATE=en_US.UTF-8
##  [5] LC_MONETARY=nl_NL.UTF-8    LC_MESSAGES=en_US.UTF-8
##  [7] LC_PAPER=nl_NL.UTF-8       LC_NAME=C
##  [9] LC_ADDRESS=C               LC_TELEPHONE=C
## [11] LC_MEASUREMENT=nl_NL.UTF-8 LC_IDENTIFICATION=C
##
## attached base packages:
```

```
## [1] stats     graphics  grDevices utils     datasets  methods   base
##
## other attached packages:
## [1] boot_1.3-25    survival_3.2-7 lme4_1.1-23    Matrix_1.2-18
##
## loaded via a namespace (and not attached):
##  [1] Rcpp_1.0.4.6    lattice_0.20-41 digest_0.6.25   MASS_7.3-53
##  [5] grid_3.6.3      nlme_3.1-149    magrittr_1.5    evaluate_0.14
##  [9] rlang_0.4.6     stringi_1.4.6   minqa_1.2.4     nloptr_1.2.2.1
## [13] rmarkdown_2.4   splines_3.6.3   statmod_1.4.34  tools_3.6.3
## [17] stringr_1.4.0   xfun_0.13       yaml_2.2.1      compiler_3.6.3
## [21] htmltools_0.4.0 knitr_1.28
```

References for the packages used follow below.

```
citation("lme4")
```

```
##
## To cite lme4 in publications use:
##
##   Douglas Bates, Martin Maechler, Ben Bolker, Steve Walker (2015).
##   Fitting Linear Mixed-Effects Models Using lme4. Journal of
##   Statistical Software, 67(1), 1-48. doi:10.18637/jss.v067.i01.
##
## A BibTeX entry for LaTeX users is
##
##   @Article{,
##     title = {Fitting Linear Mixed-Effects Models Using {lme4}},
##     author = {Douglas Bates and Martin M{\"a}chler and Ben Bolker and Steve Walker},
##     journal = {Journal of Statistical Software},
##     year = {2015},
##     volume = {67},
##     number = {1},
##     pages = {1--48},
##     doi = {10.18637/jss.v067.i01},
##   }
```

```
citation("survival")
```

```
##
## Therneau T (2020). _A Package for Survival Analysis in R_. R package
## version 3.2-7, <URL: https://CRAN.R-project.org/package=survival>.
##
## Terry M. Therneau, Patricia M. Grambsch (2000). _Modeling Survival
## Data: Extending the Cox Model_. Springer, New York. ISBN 0-387-98784-3.
##
## To see these entries in BibTeX format, use 'print(<citation>,
## bibtex=TRUE)', 'toBibtex(.)', or set
## 'options(citation.bibtex.max=999)'.
```

```
citation("boot")
```

```
##
## To cite the 'boot' package in publications use:
##
##   Angelo Canty and Brian Ripley (2020). boot: Bootstrap R (S-Plus)
##   Functions. R package version 1.3-25.
```

```
##
##   Davison, A. C. & Hinkley, D. V. (1997) Bootstrap Methods and Their
##   Applications. Cambridge University Press, Cambridge. ISBN
##   0-521-57391-2
##
## To see these entries in BibTeX format, use 'print(<citation>,
## bibtex=TRUE)', 'toBibtex(.)', or set
## 'options(citation.bibtex.max=999)'.
```
